# Overcoming software bottlenecks for scalable passive acoustic monitoring: insights from a global expert assessment

**DOI:** 10.64898/2026.03.30.715176

**Authors:** Martino E. Malerba, Cristian Pérez Granados, Kristian Bell, Maria M. Palacios, Kristen M. Bellisario, Camille Desjonquères, Alba Márquez-Rodríguez, Irene Mendoza, Christoph F. J. Meyer, Vijay Ramesh, Xavier Raick, Tessa A. Rhinehart, Connor M. Wood, Morgan A. Ziegenhorn, Giuseppa Buscaino, Marconi Campos-Cerqueira, Marina H. L. Duarte, Amandine Gasc, Tara Hanf-Dressler, Francis Juanes, Leandro Aparecido do Nascimento, Rodney A. Rountree, Karolin Thomisch, Luís Felipe Toledo, Mosikidi Toka, Manuel Vieira

## Abstract

Passive acoustic monitoring (PAM) enables non-invasive sampling of wildlife across broad spatial, temporal and taxonomic scales. Its ongoing and widespread use has generated unprecedented volumes of acoustic data, shifting the primary bottleneck from data collection to the storage, processing, integration, and interpretation of PAM outputs. Although many software tools exist to address these challenges, differences in their design, scope, and usability often create fragmented and complex analytical workflows. To identify the key barriers and opportunities shaping the implementation of PAM surveys, we conducted a structured expert solicitation involving 30 international practitioners working across terrestrial and aquatic ecosystems. Experts identified and ranked their most critical pain points in current PAM workflows, spanning data storage, processing, and interpretation. The top challenge identified related to accurate species identification using deep learning and artificial intelligence (AI) models, especially in noisy soundscapes or for underrepresented taxa. Eight additional priority challenges included workflow fragmentation, limited availability of user-friendly analytical and visualisation tools, uneven access to software, manual validation bottlenecks, computational constraints, and difficulties in data handling, standardisation, and sharing. Participants also proposed practical mitigation strategies for these priority challenges, supported by step-by-step guidance to help overcome key barriers. Together, these insights provide a roadmap toward more scalable, open-access, and collaborative software systems, which are increasingly essential to realise the full potential of PAM in global biodiversity monitoring.

## 2. Introduction

Global biodiversity is under threat, with ecosystems facing rapid declines in species richness and ecological stability (Tilman et al. 2017). Monitoring biodiversity at large scale is critical for identifying and tracking these changes, implementing conservation actions, and assessing the effectiveness of restoration efforts (Pereira and Cooper 2006). Passive Acoustic Monitoring (PAM) has become a robust and cost-effective solution for biodiversity monitoring, capturing wildlife sounds over extended periods to provide a non-invasive and scalable approach across diverse environments (Gibb et al. 2019, Darras et al. 2025). The integration of artificial intelligence (AI), and more specifically, deep learning algorithms, has further accelerated PAM adoption by enabling automated identification of target sounds in large audio datasets (Gibb et al. 2019). Yet, as PAM applications expand, so too do the challenges of storing, analysing, integrating, and interpreting the vast amount of data it generates (Gibb et al. 2019).

The affordability and performance of hardware for PAM surveys have significantly improved in recent years, marking a major milestone for its scalability (Sheng et al. 2019). Early adoption of PAM relied on a limited choice of commercial recorders, which, although reliable, had a cost exceeding USD 1,000 per terrestrial unit and USD 5,000 per aquatic unit (REF here). These costs constrained adoption, particularly in resource-limited contexts and large-scale monitoring programs. The emergence of lower-cost devices such as AudioMoth (Hill et al. 2018) and Song Meter Micro (Wildlife Acoustics, USA) has increased accessibility for PAM, particularly for terrestrial applications, with unit costs now commonly around USD 100-150 (Gibb et al. 2018). This hardware transition enables unprecedented data collection across large spatial and temporal scales (Sugai et al. 2019, Stowell and Sueur 2020, Hoefer et al. 2023). As a result, the primary bottleneck in PAM has shifted from data acquisition to data management, analysis, and interpretation (Gibb et al. 2019).

In parallel with rapid advances in PAM hardware, the quantity, complexity, and diversity of acoustic data being generated have increased dramatically. While major progress has been made in automated detection and classification, particularly through the rise of deep-learning and foundation bioacoustic models (Kahl et al. 2021, Hamer et al. 2023, van Merriënboer et al. 2025), these advances have not been matched by equivalent progress in the broader software ecosystems required to manage, analyse, and interpret large PAM datasets. In practice, PAM users must navigate a fragmented landscape of tools for data storage, metadata integration, species detection, manual validation, visualisation, and reporting, often across different programming languages, file formats, and platforms (Hanf-Dressler et al. 2026). Although some web-based platforms now offer cloud storage and collaborative features (e.g., Aide et al. 2013, Darras et al. 2024), many workflows still rely on *ad hoc* combinations of software that are difficult to scale, transfer, and maintain across projects and user groups. As a result, despite clear technological progress, software-related barriers continue to limit the efficiency, reproducibility, and widespread adoption of PAM in biodiversity monitoring.

In this study, we employed a structured expert consultation to identify the key technical and practical barriers limiting the scalability, accessibility, and reproducibility of PAM analyses for biodiversity monitoring. Open-ended responses were scored and ranked to prioritise the pain points perceived as most limiting by practitioners. We first synthesise the dominant challenges reported by the community, spanning species identification, fragmented analytical workflows, data handling, validation, and standardisation. We then summarise expert-endorsed strategies currently used to mitigate these barriers, highlighting practical tools and approaches already available. We also translate these insights into a set of practical guides provided as Supplementary Material to support implementation in real-world PAM workflows (see Box 1). Finally, we outline how emerging developments, such as edge computing, cloud-based processing, and citizen science integration, can offer transformative solutions to scale PAM globally. Together, these insights provide a roadmap for turning PAM into a more accessible, interoperable, and impactful tool for biodiversity monitoring and conservation.

### Box 1.

**Practical guides provided as Supplementary Material.**

To translate the outcomes of the expert consultation into actionable resources, we provide a series of practical guides as Supplementary Material (Guides). Each guide addresses a specific software or workflow challenge commonly encountered in large-scale passive acoustic monitoring (PAM) and illustrates feasible solutions using currently available tools and approaches.

Guide 1. *Expanding BirdNET models to non-avian species through transfer learning*: Using a case study of a multi-species frog classifier, it demonstrates how an existing CNN platform can be repurposed to detect non-avian taxa using transfer learning, enabling the development of custom classifiers from relatively small, taxon-specific training datasets.

Guide 2. *Building acoustic classifiers with little or no labelled training data*: Introduces embedding-based and clustering approaches that allow researchers to discover, curate, and label recurrent sound types directly from unlabelled recordings, supporting classifier development in data-poor systems.

Guide 3. *Data augmentation in PAM workflows*: Reviews dataset-level and signal-level data augmentation strategies that improve classifier robustness and generalisation, with practical guidance on when and how to apply augmentation in bioacoustic contexts.

Guide 4. *Coding an integrated multi-tool workflow in R*: Provides a step-by-step framework for building reproducible, end-to-end PAM pipelines in R, integrating audio handling, detection, classification, validation, and visualisation across multiple tools.

Guide 5. *Developing GUI-based interfaces for PAM*: Explores how graphical user interfaces can lower technical barriers to PAM by supporting tasks such as validation, annotation, threshold selection, and exploratory analysis without requiring coding expertise.

Guide 6. *Accounting for classifier errors to make correct ecological inferences*: Presents statistical and validation-based strategies for managing false positives and false negatives in automated detector outputs, ensuring that ecological inferences remain robust and defensible.

Guide 7. *Using metadata standards for acoustic data management*: Outlines best practices for structuring and documenting PAM datasets using hierarchical metadata, drawing on existing biodiversity standards to improve data interoperability, reuse, and long-term value.

Guide 8. *Reproducible scientific workflows for PAM*: Describes practical approaches to ensuring transparency and reproducibility in PAM research, with a focus on version control, documentation, open-source tools, and collaborative platforms such as Git and GitHub.

## 3. Material and Methods

### 3.1. Expert Identification

We compiled an initial list of PAM specialists by selecting the top 100 most-cited researchers in the field (each with a minimum of five peer-reviewed publications on this topic). Citation data were extracted from the Web of Science platform on the 4^th^ of June 2025. This list was subsequently expanded through purposive and snowball sampling, incorporating additional experts identified through collaborator recommendations and their recognised roles in prominent PAM projects, monitoring programs, and software development initiatives. The combined approach ensured representation across sectors, geographic regions, ecosystem types, and taxonomic expertise. Our final invitation list comprised 156 experts, who were contacted via email by MEM and CPG between 9^th^ and 13^th^ June 2025. To encourage participation, two reminder emails were issued during the survey period.

### 3.2. Initial expert consultation: Identifying and Prioritising Pain Points

We conducted an expert consultation through an online semi-structured questionnaire comprising 10 questions (see Supplementary Material Questionnaires), designed to capture both quantitative and qualitative data on current practices and challenges in PAM. The questionnaire was structured into three components:

- **Background information:** Captured participants’ domain expertise (e.g., birds, marine mammals), years of experience, geographic and environmental focus, and professional role.
- **Hardware and software usage:** Collected data on the types of PAM hardware and software used, including specific tools for species identification, metadata management, and AI model deployment.
- **Pain points:** Respondents were asked to write (max one sentence length) and rank in order of importance their top five challenges for processing PAM data, including storage, annotation, analyses, and validation.

The survey instrument was developed in the Qualtrics platform and piloted internally by MEM, MP, and CPG to ensure clarity and relevance. It was administered online and distributed via email between 9^th^ and 25^th^ of June 2025. Participation was voluntary, and responses were anonymous unless participants opted to provide contact details for follow-up engagement or co-authorship in subsequent outputs. Participants were able to skip questions, resulting in variation in response numbers across items. A total of 42 respondents completed the survey, with responses per question ranging from 33 to 42.

### 3.3. Analysis and prioritisation of Pain Points

Quantitative and qualitative analyses were undertaken to identify, synthesise, and prioritise key challenges in PAM workflows. Qualitative responses to open-ended questions were first analysed using an inductive thematic approach. In total, nine challenges were reported across all experts and subsequently synthesised through an iterative coding process. Challenges were grouped based on conceptual similarity, with overlapping items merged and terminology standardised. Coding and categorisation were conducted by MEM and refined through discussion until agreement was reached, resulting in a consolidated set of nine core pain points.

This synthesised set was then shared with all 42 PAM experts via email in August 2025 for validation, ensuring completeness and accurate representation of the issues identified. Following validation, quantitative ranking data provided by participants were aggregated to generate a total severity score for each pain point, enabling comparison of relative importance and identification of the most critical bottlenecks in PAM workflows.

### 3.4. Follow-up expert consultation: Strategies to Minimise Pain Points

We developed and distributed a second survey instrument to identify practical strategies for addressing the challenges identified in the initial consultation. The online Qualtrics survey was distributed via email by CPG to the same pool of 42 PAM participants in August 2025, with 24 experts completing the questionnaire.

The survey instrument comprised open-ended questions designed to elicit both current practices and future-oriented solutions. To minimise ordering bias, the list of identified pain points was randomised for each participant. For each pain point, experts were asked to: (1) describe existing tools, methods, and practices currently used to address the issue, and (2) outline what an ideal, yet realistic, future solution would look like.

Responses were analysed using an iterative qualitative synthesis approach. The core team reviewed and consolidated responses by grouping similar strategies, standardising terminology, and removing redundancies. The resulting synthesis was then circulated to the 24 PAM experts who completed the second survey via email on 20^th^ October 2025 for validation, refinement, and identification of any gaps or overlooked areas. Through this iterative feedback process, we reached consensus on a final set of recommended strategies to address each pain point, including both currently available solutions and priority areas for further development.

All co-authors reviewed and approved the final list of pain points and proposed solutions, ensuring shared ownership and alignment with community needs. For each pain point, we document at least one illustrative case study of existing practices or tools to address that challenge (see Box 1).

## 4. Results

### 4.1. Demographic Overview of Survey Respondents

Overall, we received complete sets of answers from 30 PAM experts. Most experts were academic researchers (87%), with the remaining from industry or from government agencies. The majority of experts had >8 years of experience (80%), followed by 4-7 years (14%) and 1-3 years (6%).

In terms of geographic region where the experts have conducted work, the survey captured input from all continents (Fig. 1). Most respondents have collected data in Europe (31%), South America (25%), and North America (22%), following up were Africa (17%), Asia (12%), Australia & Oceania (11%) and Polar Regions (8%) (Fig. 1A). Across realms, a larger number of experts worked mainly in terrestrial ecosystems (55%), followed by marine environments (30%) and freshwater systems (15%; Fig. 1B).

**Figure 1:**
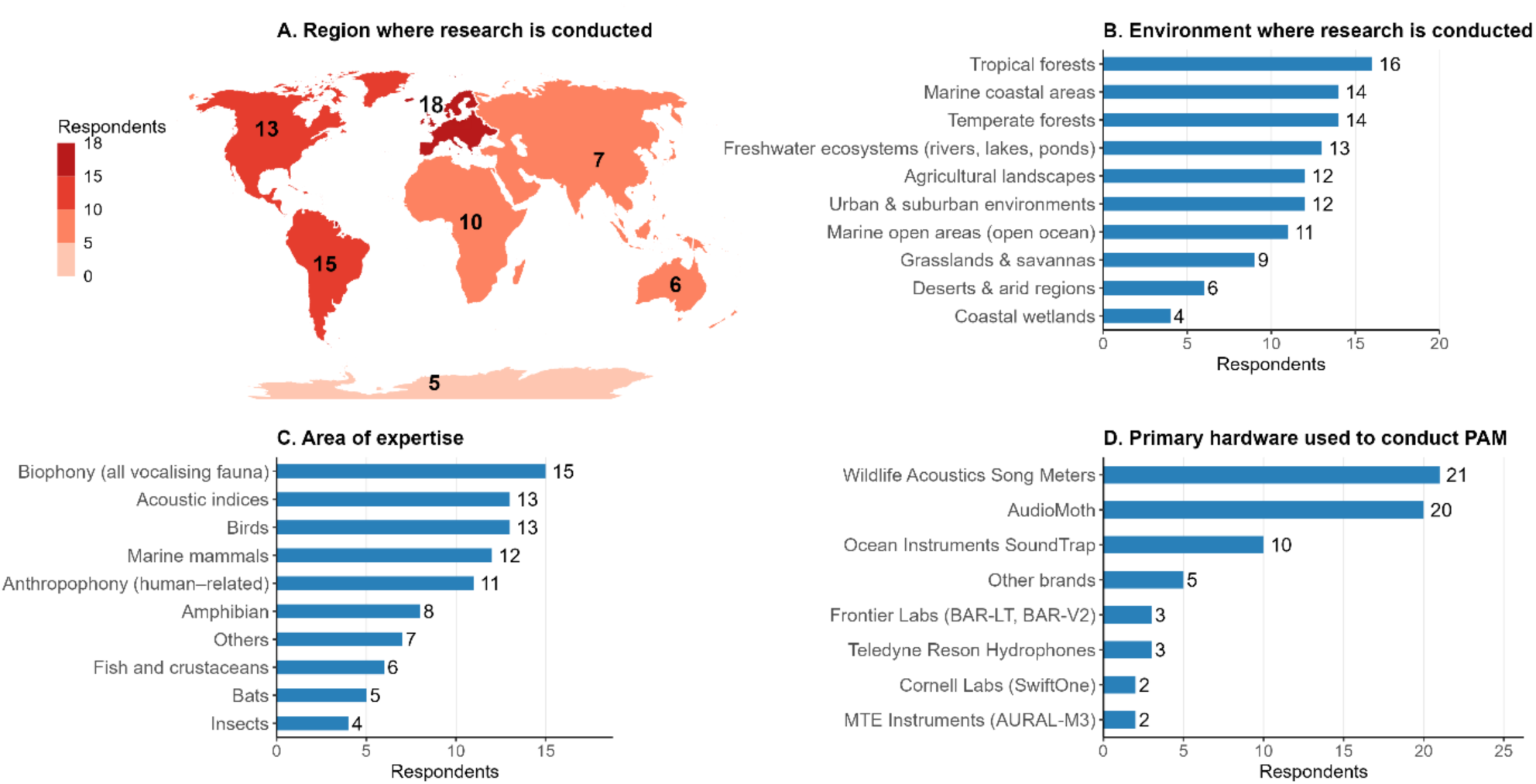
Geographic regions (A), research environments (B), areas of expertise (C) and hardware types used (D) reported by survey respondents (n = 42). Values indicate the number of respondents selecting each category, and respondents could select multiple options.

**Figure 2:**
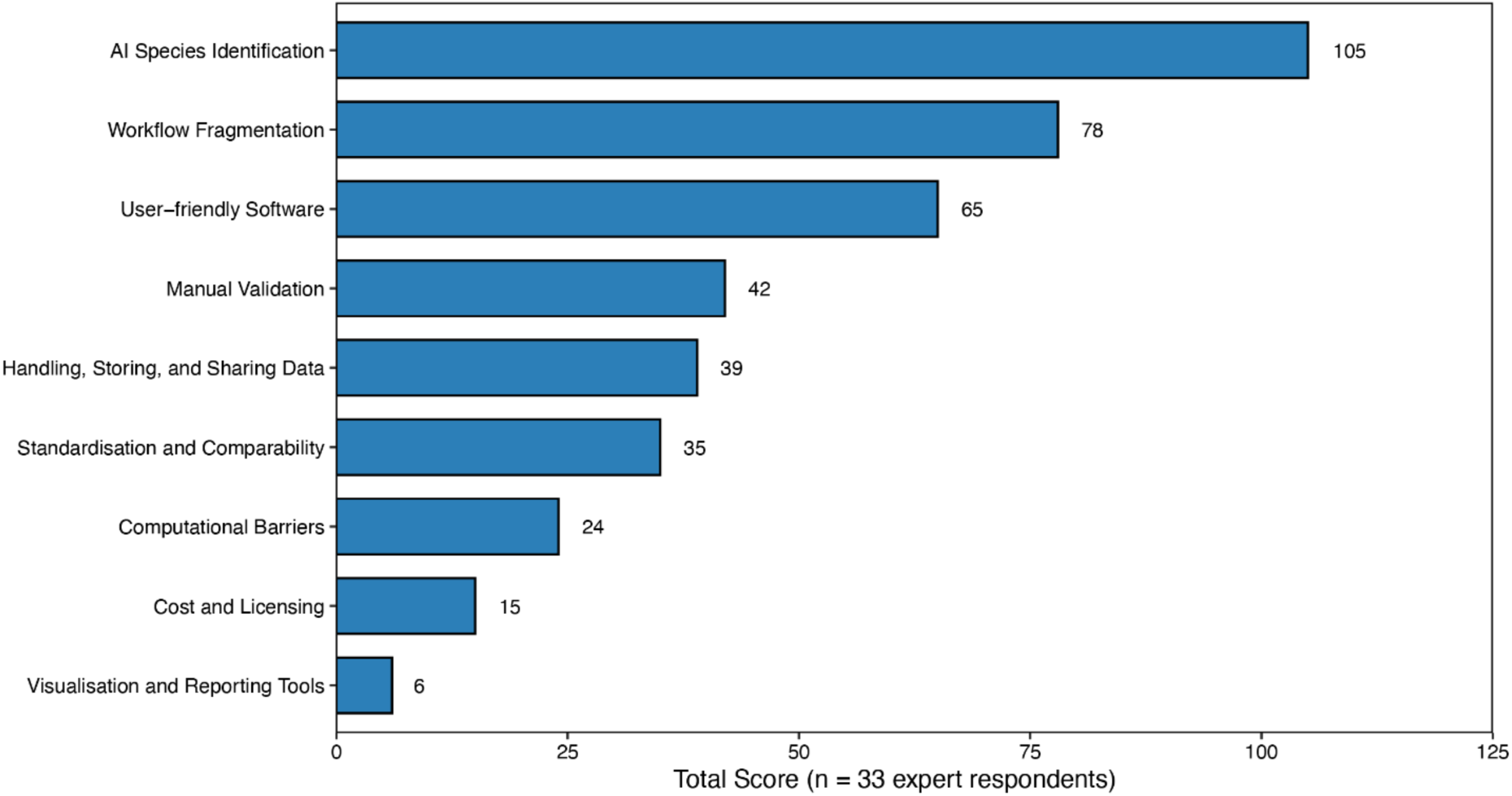
Total perceived importance of key pain points in passive acoustic monitoring (PAM) workflows, ranked by expert respondents. Each bar represents the cumulative score for a broad category of challenges, weighted from 5 (most important) to 1 (least important) according to respondents’ rankings. Categories are ordered by total score from highest to lowest.

Expertise was most frequently reported for biophony (15.9%), avian monitoring (13.8%), and marine mammal monitoring (12.8%), amphibians (8.5%), fish and crustaceans (6.4%), bats (5.3%), and insects (4.3%; Fig. 1C). Several respondents also reported experience with terrestrial mammals, including primates and large carnivores (2.1%). Beyond taxon-specific applications, a substantial proportion of expertise is related to soundscape and analytical domains, including acoustic indices (13.8%), anthropophony (11.7%), and geophony (3.2%). Finally, two respondents identified themselves as software developers and taxonomic bioacousticians (Fig.1C).

When it comes to PAM hardware, Wildlife Acoustics Song Meters (33%) and AudioMoth devices (31%) emerged as the most widely used (Fig. 1D). Other devices cited were the Ocean Instruments SoundTraps (16%), for underwater applications, followed by devices from Frontier Labs, Teledyne Reson, MTE Instruments, or SwiftOne (2–3 respondents each). A few participants reported alternative systems, including SNAP Recorders (Loggerhead Instruments), MARU and Rockhopper recorders (K. Lisa Yang Center for Conservation Bioacoustics), Cetacean Research Technology, and SonoVault recorders (Fig. 1D).

### 4.2. Pain points for analysing PAM data

The experts identified nine overarching themes derived from 122 individual text entries describing key pain points in PAM workflows (see *Supplementary Material Questionnaires* for categorisation of text responses). *AI-based species identification* emerged as the most critical challenge, receiving the highest cumulative score (105), reflecting widespread concern about the limited accuracy, transparency, and cross-taxon applicability of current AI classification models. The second most prominent theme, *workflow fragmentation* (78), underscored frustration with disconnected tools, and the need for integrated analysis pipelines. The lack of u*ser-friendly software* ranked third (65), with many respondents emphasising steep learning curves and limited accessibility of current platforms. Other recurring themes included *manual validation* (42), *data handling and sharing* (39), and *standardisation and comparability* (35), all pointing to barriers that limit efficiency and reproducibility. Lower-ranked but still notable themes included *computational barriers* (24), *cost and licensing* (15), and *visualisation and reporting tools* (6).

Despite variation in scores, all themes included answers from multiple experts, indicating a broad and interrelated set of challenges currently limiting the scalability and adoption of PAM.

#### 4.2.1. AI Species Identification (Total Score: 105 – Highest Priority)

Improving how we use deep learning models and AI more broadly to identify species accurately and efficiently was voted as the most urgent challenge in PAM. Across ecosystems, PAM is hampered by error rates in automated species identification (Barré et al. 2019, Metcalf et al. 2022). The performance of AI models greatly varies across taxa, regions, recorders, and recording conditions, especially in the presence of dialect variation, intra-specific variability, or species with similar calls, as well as background noise in aquatic recordings (Gibb et al. 2019, Hall et al. 2025). Therefore, there is a need for robust and standardised validation procedures to avoid misinterpretation of outputs. Another recurring issue identified in the survey is the lack of open-access annotated reference datasets to train or benchmark models, as annotating is a time-demanding process that requires great expertise in the studied species. In the absence of open, standardised calibration datasets and geographically diverse training data, even models that perform well under benchmark conditions can fail unpredictably when applied to new species, regions, or recording setups, limiting their reliability and transferability in real-world PAM applications.

##### 4.2.1.1. Current Strategies in Use

A widely endorsed approach for improving AI-based species identification in real-world PAM workflows involves customising existing foundation models or pretrained architectures (e.g., BirdNET and Perch) to new taxa or acoustic contexts using transfer learning, either within or across domains (e.g., Bell et al. 2026, Ghani et al. 2023; Huang et al. 2022, Márquez-Rodríguez et al. 2026, Williams et al. 2025). This strategy allows researchers to leverage the robustness of established models while addressing persistent gaps in taxonomic and regional coverage, including poorly represented taxa (see SI Guide 1: *Expanding BirdNET models to non-avian species through transfer learning*) and situations where labelled training data are scarce or unavailable using model embeddings and clustering approaches (see SI Guide 2: *Building acoustic classifiers with little or no labelled training data*). To further mitigate the scarcity of annotated data, particularly for rare or understudied species, several experts highlighted the use of data augmentation techniques to artificially expand training datasets (see SI Guide 3: *Data augmentation in PAM workflows*).

Experts also consistently emphasized that automated detection alone is insufficient for reliable inference, underscoring the need for workflows that combine AI-based classification with targeted manual validation (see SI Guide 6. *Accounting for classifier errors to make correct ecological inferences*). Practical strategies included prioritising subsets of detections for expert review, applying biologically informed thresholds and statistical filters (e.g., Knight et al. 2017, Wood and Kahl 2024, Wood and Peery 2022), and using ensemble approaches that combine outputs from multiple models to reduce error rates (Pérez-Granados et al. 2023). Collaboration emerged as a cross-cutting solution. Respondents advocated pooling annotated datasets across projects and regions, engaging citizen science to expand validation capacity or refine existing models, and sharing open code and workflows to support community-driven improvements.

These needs were seen as particularly acute in aquatic environments, where most recorded sounds remain unidentified, motivating collaborative initiatives such as FishSounds (Looby et al. 2022) and the Global Library of Underwater Biological Sounds (GLUBS; Parsons et al. 2024).

##### 4.2.1.2. Emerging Solutions

Looking ahead, respondents converged on the need for globally diverse, open-access, and reliable annotated datasets, as well as generalisable AI models capable of handling a wide range of taxa, geographies, and recording conditions. Several envisioned a collaborative platform that would function as both a model- and dataset-sharing hub, enabling researchers to upload, adapt, and retrain models in a streamlined way. The development of explainable AI (tools that make model decisions interpretable) was highlighted as critical for building user trust. Many experts also called for built-in uncertainty metrics and user-friendly interfaces that would allow ecologists without coding skills to confidently assess and adapt outputs. Some stressed the value of foundational AI models, akin to large language models, that could be fine-tuned for specific ecological contexts or taxa. Others saw the future in standardised protocols for training and validation, ensuring comparability across studies. Although fully generalisable “all-species” detectors may be aspirational, respondents saw clear potential in scalable frameworks that integrate local adaptation, ensemble modelling, and transparent reporting, allowing AI species identification to mature into a robust, reliable tool for global biodiversity monitoring.

#### 4.2.2. Workflow Fragmentation (Total Score: 78)

Respondents highlighted that workflows regarding PAM processing remain fragmented. Navigating from data storage to acoustic analyses, model validation, statistical analyses and data reporting often requires using several different tools, which often lack compatibility. For example, the detector output from one software may not integrate seamlessly with manual annotation platforms or downstream statistical packages. This creates inefficiencies, increases the risk of data loss or misinterpretation, and makes it harder to scale up analyses across large datasets or multiple projects.

##### 4.2.2.1. Current Strategies to Manage Workflow Fragmentation

Respondents highlighted that the main way to cope with workflow fragmentation today is by manually stitching together disconnected tools while trying to maintain consistency through careful file organisation, documentation, and coding. Many rely on R and Python pipelines (e.g., warbleR, vegan, monitoR, or seewave for R and scikit maad for Python) to integrate outputs from multiple programs (see SI Guide 4: *Coding an integrated multi-tool workflow in R*), while others use modular platforms like Arbimon, PAMGuard, Raven, or EcoSoundWeb to cover as many steps as possible in one place. A recurring strategy is to enforce standardised file formats (e.g., CSV, metadata schemas) to reduce incompatibilities, and to adopt relational databases or dashboards for centralised visualisation and reporting. Users also mentioned creating custom templates and Standard Operating Procedures (SOPs) to maintain reproducibility across teams, while some collaborate directly with software developers to encourage compatibility. Despite these strategies, respondents acknowledged that fragmentation often still requires substantial manual work, and that maintaining pipelines remains time-consuming.

##### 4.2.2.2. Emerging Solutions

The vision for an ideal solution was clear: a fully integrated, interoperable, and modular PAM platform that seamlessly handles annotation, detection, classification, validation, metadata management, storage, and reporting within a single workflow. Some envisioned a “one-stop shop” solution akin to ArcGIS or QGIS for acoustics, where all modules interconnect but can also be updated independently as methods evolve. Several suggested a cloud-based infrastructure, offering scalability for large datasets, automated documentation of workflows, and real-time collaboration across teams. Respondents emphasized that such a platform should adopt universal standards for annotation formats, acoustic features, and metadata, while remaining flexible enough for local adaptation. Importantly, respondents emphasized that the future platform must be user-friendly and either open source or have commercial licensing models reserved exclusively for corporate use. Since completion of the survey, development toward such a modular PAM platform has begun within the Raven environment. However, key aspects remain absent or unclear, including cost of access, support for multi-user collaboration, the ability to search similarity among embeddings, and integrated data storage capabilities.

#### 4.2.3. User-Friendly Software (Total Score: 65)

A major barrier to wider use and innovation in PAM is the technical complexity of the tools. Many researchers and practitioners in the fields of ecology or applied conservation are not professional programmers, which often reduces the potential of citizen science in PAM surveys. Although several current platforms already provide graphical user interfaces (GUIs; see the next section), a large fraction of tools available for PAM processing still do not offer such advantages and require advanced coding skills or familiarity with programming environments such as R, Matlab or Python (see Rhinehart et al. 2024 for an overview of available bioacoustics software).

##### 4.2.3.1. Current Strategies

Researchers often rely on GUI-based tools (e.g., BirdNET, Kaleidoscope, Raven, Arbimon, PAMGuard, Triton, and WildTrax). However, each has limitations in taxonomic coverage, transparency, or accessibility due to cost. A common strategy is to combine these GUI tools with tutorials, training workshops, and shared annotated R/Python scripts (e.g., R Markdown for R and Jupyter notebooks for Python) to help non-programmers bridge gaps. Some researchers reported developing their own Shiny Apps or coding add-ons to provide step-by-step guidance (see SI Guide 5: *Developing GUI-based interfaces for PAM*), while others highlighted collaboration with computer scientists as an effective way to access more advanced tools. Another recurring theme was the need for clear documentation, reproducible guides, and community support to make existing software more approachable for ecologists and managers with limited coding experience.

##### 4.2.3.2. Emerging Solutions

Looking ahead, respondents consistently envisioned a more inclusive PAM landscape where user-friendly software is the norm. The ideal solutions would be open-access, GUI-based platforms capable of guiding users through every step of the workflow without requiring coding: detection, classification, validation, and reporting. Such platforms should be modular, intuitive, interoperable, and adaptable across taxa and environments, with consistent terminology, integrated tutorials, and context-sensitive help. Others suggested that collaboration with big tech companies or startups could accelerate the development of robust, affordable tools, potentially integrating co-pilot features to support occasional coding needs. Ultimately, respondents hope for software ecosystems that shift the focus away from technical hurdles and back to ecological interpretation and conservation impact.

#### 4.2.4. Manual Validation (Total Score: 42)

Manual review of model predictions is essential for ensuring reliability and ecological validity of automated outputs. However, respondents highlighted a lack of software tools that support efficient, structured, and transparent validation, including built-in quality controls, prioritisation of detections (e.g., by confidence scores), or clustering of acoustically similar calls to streamline review. As a result, current workflows often require users to switch between multiple, poorly integrated tools such as detection software, spectrogram viewers, and spreadsheets, increasing both time costs and the risk of error. There is therefore a strong need for integrated validation environments that preserve the critical role of expert judgement while making manual review faster, more consistent, and more collaborative.

##### 4.2.4.1. Current Strategies

To cope with the heavy burden of manual validation, respondents emphasized combining automation with selective human oversight. Many use tools such as BirdNET, Raven Pro, Triton, DetEdit, Arbimon, and Kaleidoscope, often integrating them with custom scripts to streamline spot-checking and label editing. A recurring strategy is to validate only subsets of detections, prioritising uncertain or low-confidence outputs, or using statistical models (e.g., occupancy models) to account for detection errors instead of exhaustively verifying every prediction (see SI Guide 6: *Accounting for classifier errors to make correct ecological inferences*). Others recommended clear annotation guidelines, training multiple users to share the load, or employing consensus approaches to reduce subjectivity. Several also highlighted the importance of open-access, pre-validated datasets that allow users to benchmark models and minimise repetitive validation work (see Pérez-Granados et al. 2025). Active learning and filtering approaches (e.g., focusing on species known to occur at certain locations or start reviewing high-probability predictions for presence/absence) are practices currently implemented to reduce time and effort without losing reliability, as well as citizen science initiatives (e.g., Forest Listeners project; Google Arts & Culture n.d.).

##### 4.2.4.2. Emerging Solutions

Experts envisioned semi-automated, AI-assisted validation systems that minimise human workload while maintaining transparency and trust in the validation process. Desired features include intuitive review interfaces, built-in confidence scoring, consensus tools, and error metrics that help prioritise ambiguous detections. Consensus emerged for centralised, open-access repositories of annotated datasets that could serve as benchmarks, training resources, and validation references across taxa and regions. Many also expressed interests in cloud-based platforms with integrated detection, validation, and statistical tools, enabling collaborative review and large-scale consistency. In an ideal scenario, detection models would become accurate enough that manual validation is rarely required, with human review reserved primarily for quality control or rare or unknown sound types; however, this outcome is unlikely to be realised in the near term. As such, agile interfaces that combine automated detection, guided human annotation, and continuous model improvement were viewed as the most promising path forward.

#### 4.2.5. Handling, Storing, and Sharing Data (Total Score: 39)

Data management challenges are growing as more recorders are deployed over longer periods. Some teams now work with petabytes of data, yet storage infrastructure, backup solutions, sound archive infrastructure, and metadata management have not kept pace. File formats, although most projects use .WAV, are often inconsistent across devices or projects, and few tools offer built-in metadata validation or compatibility with online biodiversity repositories like iNaturalist, Xeno-Canto or GBIF. Sharing data across institutions, sound archives, or even among team members can be painful due to access restrictions, lack of cloud integration, or poor searchability within archives. There is a lack of standard protocols for organising folders, naming files, or synchronising annotations, which greatly reduces the degree of interoperability among projects. These issues reduce reproducibility, increase overhead, and make it hard to contribute to broader meta-analyses or cumulative research efforts.

##### 4.2.5.1. Current Strategies

Respondents described a mix of stop-gap solutions for managing and sharing large acoustic datasets. Many rely on external hard drives, SSDs, NAS systems, and fragmented cloud storage (e.g., Box, Google Drive, Amazon Web Services, Arbimon, WildTrax) or repositories (e.g., Zenodo) to store and transfer data. For some teams, a common workaround is to use compressed, open formats (e.g., FLAC), although most use original, non-compressed formats (e.g., .WAV). Some emphasised the importance of adopting metadata standards, including file naming, (e.g., Darwin Core (Wieczorek et al. 2012), GUANO (Riggs, 2018) or Tethys (Roch et al. 2013),) to improve interoperability (SI Guide 7: *Using metadata standards for acoustic data management*). Overall, strategies remain fragmented and resource-dependent, with special difficulties in the Global South where backup capacity is limited.

##### 4.2.5.2. Emerging Solutions

Although depositing sound files in well-established sound archives is still a recommended and incentivized practice (Toledo et al. 2015, Dena et al. 2018, Mendoza-Henao et al. 2021), experts widely agreed on the need for a centralised, open-access repository for PAM data, ideally with standardised metadata requirements and globally agreed file formats. Many envisioned a cloud-based system (affordable, scalable, and user-friendly) that would automate backups, metadata capture, provenance tracking, and collaborative sharing. Some saw potential in building a global network of regional data nodes to distribute storage costs and ensure long-term preservation. In parallel, respondents called for stronger investment in high-biodiversity regions and free storage for conservation-focused projects, supported by governments, institutions, or large technology providers. A more realistic near-term goal may be improved interoperability between existing archives and analysis platforms, enabling data to be stored, discovered, and reused without being lost at the end of individual projects.

#### 4.2.6. Standardisation and Comparability (Total Score: 35)

Despite the diversity of tools and platforms for processing PAM data (e.g. Hanf-Dressler et al. 2026), there are few guidelines and standards about what constitutes best practice. This hinders the ability to compare results across projects or build general knowledge. Metrics such as signal-to-noise ratio, detection thresholds, or classification accuracy are often reported inconsistently or omitted entirely, with few tools including calibration routines or reference levels to allow for instrument comparability. Major differences also exist between terrestrial and aquatic research, reflecting the distinct characteristics and monitoring requirements of these environments. In addition, required metadata fields, and their structure, differ widely between software tools, and users commonly struggle to merge datasets without extensive cleaning. What is needed are community-driven standards for data formats, analysis parameters, and reporting structures, perhaps with minimal but enforceable core requirements to ensure interoperability and data longevity.

##### 4.2.6.1. Current Strategies

Respondents highlighted that the lack of standardisation is a persistent barrier but suggested practical steps to improve comparability. Many stressed the importance of precise reporting of study design, hardware, and software parameters to allow further replicability and comparisons, following the path of MANTA (“Making Ambient Noise Trends Accesible”, Miksis-Olds et al. 2021). Some recommend adopting core sets of metrics that can be consistently reported across projects, while leaving flexibility for study-specific needs. Such adoption could start with current existing metadata standards (e.g., Darwin Core, ISO initiatives, Acoustical Society of America guidelines; see SI Guide 7: “*Using metadata standards for acoustic data management*”) and develop best-practice protocols that are easy to follow. However, this is further complicated by requirements that often vary across regions and realms. Implementing collaborative approaches, such as workshops, international networks, and cross-project benchmarking exercises, is an effective tool for promoting transparency and alignment across studies (see e.g. Wall et al. 2025).

##### 4.2.6.2. Emerging Solutions

Experts envisioned the creation of ISO-level guidelines or open-access handbooks with clear, taxa-specific recommendations, supported by calibration protocols and benchmarking datasets. Ensuring dedicated funding and global consortia to drive standardisation efforts and aligning ecoacoustics with existing frameworks (e.g., GOOS (Tyack et al. 2023), IQOE (Tyack et al. 2015), TG Noise (Van der Graaf et al 2012)) will contribute to the required standardisation. There was broad consensus that an open, published international standard, broken down by group of taxa, environments, or regions, would greatly improve reproducibility and ecological synthesis, enabling PAM outputs to be compared meaningfully across scales.

#### 4.2.7. Computational Barriers (Total Score: 24)

Although computational resources have improved, many PAM users still encounter bottlenecks, especially when processing large amounts of audio or using custom deep-learning models. Training or fine-tuning models often requires GPUs or cloud infrastructure that may be unavailable or expensive. Even seemingly basic steps like spectrogram loading or batch processing can become slow on older systems or when working remotely. These delays are particularly acute in projects aiming for real-time or near-real-time analysis. Without better tools for parallel processing, optimised storage formats, or lightweight review interfaces, PAM is unlikely to achieve the level of scalability required for large-scale biodiversity monitoring, at least for certain groups and regions.

##### 4.2.7.1. Current Strategies

Respondents described a range of pragmatic and collaborative strategies to overcome computational barriers in PAM analyses. Experts commonly rely on university high-performance computing clusters, public servers, or cloud-based platforms (e.g. Google Cloud, AWS, Arbimon, EcoSoundWeb), while others partner with colleagues or institutions that have access to advanced computing infrastructure. To reduce computational demands, experts reported optimising workflows through data subsampling, file compression, and the use of foundation models such as BirdNET and Perch, including fine-tuning existing models for specific applications. Several respondents also emphasised the importance of realistically planning PAM data collection to match available computational resources, as well as integrating parallel processing and lightweight R or Python pipelines to maximise efficiency on standard desktop or laptop computers.

##### 4.2.7.2. Emerging Solutions

Looking ahead, experts envisioned user-friendly, cloud-based platforms with equitable access to computational resources as the long-term goal for supporting PAM workflows at scale. Ideally, these platforms would provide free or low-cost access, particularly for researchers and conservation groups in the Global South, while integrating parallel processing, containerisation, and built-in optimisation to handle very large datasets efficiently. Some respondents also pointed to the potential of shared or distributed computing approaches, in which idle processing capacity could be pooled across institutions, drawing inspiration from citizen science models such as SETI@home (Anderson et al. 2002). However, no current platform fully addresses these needs. Existing solutions such as Arbimon offer several elements of this approach, but uncertainty over continued free access and limited cross-platform flexibility in the classifiers that can be applied deter some users from full adoption. As such, the broader vision of fully integrated, cloud-ready PAM platforms capable of handling very large datasets efficiently remains largely unrealised.

#### 4.2.8. Cost and Licensing (Total Score: 15)

Cost and licensing constraints remain a significant barrier to the uptake and scalability of PAM workflows, particularly for researchers with limited funding or institutional support. Many widely used PAM and bioacoustics platforms rely on proprietary licenses, with annual fees typically ranging from hundreds to several thousand USD per user or project, depending on functionality and data volume, which can restrict access to advanced features or limit team-wide use. In addition, cloud-based processing introduces recurring costs for storage, computing, and data transfer, which can quickly accumulate to hundreds or thousands of USD per year for projects managing large acoustic libraries or supporting collaborative access. Together, these costs can force reliance on outdated or poorly maintained free tools. In this context, open-source software with permissive licenses and transparent infrastructure costs is essential for ensuring equitable access to PAM tools and for supporting reproducible and globally inclusive methodological innovation.

##### 4.2.8.1. Current Strategies

Respondents emphasised that many effective free or open-source tools are already available, such as BirdNET, Arbimon, Perch, PAMGuard, PamGUIDE, MANTA, RavenLite, R, and Python packages. Where proprietary tools exist (e.g., Kaleidoscope Pro, Raven Pro, Matlab), researchers often rely on free versions, discounted licenses, or region-specific exemptions (e.g., Raven Pro licenses for researchers in the Global South). Many also favor open-source workflows by coding in R or Python, creating their own scripts, or sharing tools via GitHub, Zenodo or other public repositories (SI Guide 10: *Reproducible scientific workflows for PAM*). Collaborative approaches, such as negotiating team-wide licenses, forming networks to apply for funding, or using institutional access, were also suggested as ways to reduce individual costs and ensure broader accessibility. Overall, there is a strong culture of working with and promoting free, open alternatives wherever possible. Experts envision a future where all PAM processing tools are free, open-source, and sustainably supported, ensuring equity across research communities worldwide. While this remains largely aspirational at present, many responses called for long-term, publicly funded support models to maintain and update free tools, with possible tiered systems where companies pay licenses but non-profit and academic researchers have free access. Others stressed the importance of building a unified, open-source PAM platform that is modular, user-friendly, and well-maintained, rather than fragmented tools with limited funding. Suggestions included government or philanthropic sponsorship, collaborative funding networks, and infrastructure to ensure transparency and reusability. The overarching goal is to remove licensing barriers, promote global equity, and accelerate collaboration and innovation in PAM.

#### 4.2.9. Visualisation and Reporting Tools (Total Score: 6 – Lowest Priority)

While not the top concern, experts agreed that it is challenging to turn PAM data into clear, actionable insights. Many tools lack effective visualisation options, such as to display presence/absence maps, species trends, or annotated spectrograms for review. Reporting functionalities are also limited, requiring manual compilation of figures or metrics for funders, managers, or collaborators. Such reporting should always include a description of the parameters used for recording and spectrogram visualisation (i.e. FFT size, overlap, frequency resolution) to ensure reproducibility across studies, an aspect emphasised by several respondents. Better integration with camera-trap outputs, environmental metadata, and mobile-friendly platforms would enhance usability and communication. While this category had the lowest total score, its identification as a pain point highlights its importance in supporting uptake and outreach beyond the research community.

##### 4.2.9.1. Current Strategies

To overcome gaps in visualisation and reporting, many experts reported relying on a mix of tools (e.g., Audacity, Raven, BirdNET-Analyser GUI, PAMGuard, Triton) and custom R and Python scripts using open-source packages (e.g., seewave, warbleR, monitoR). Business intelligence software (e.g., Power BI) and R Shiny dashboards are also emerging as ways to create interactive, shareable reports (see SI Guide 5: *Developing GUI-based interfaces for PAM*). Integrating PAM outputs with environmental metadata, eDNA, or camera trap data is challenging (Lombardo et al. 2026), and currently requires bespoke coding solutions. Collaborative platforms like Arbimon and WildTrax provide partial solutions by linking PAM with other ecological data, but fragmentation remains.

##### 4.2.9.2. Emerging Solutions

Looking ahead, the vision converges on the development of integrated and user-friendly platforms capable of combining metadata integration, detection, validation, visualisation, and reporting in one streamlined workflow (Dantzker et al 2025, Rountree et al. 2020). Respondents imagined systems that could automatically generate interactive spectrograms, diel or seasonal summaries of the predictions validated, and stakeholder-ready reports without requiring extensive coding in R or Python. Such platforms should allow seamless incorporation of environmental variables, GPS, eDNA, and camera trap data in a spatially explicit way, enabling ecological inference such as occupancy or density models. Automated quality checks for issues like clipping or internal noise were seen as valuable additions to accelerate early-stage validation. Importantly, these systems should be open-source, modular, and reproducible, evolving alongside community needs. The overarching ideal resembles a unified ecosystem (something akin to combining Arbimon, Raven, Shiny dashboards, QGIS, and eBird) capable of serving both researchers seeking detailed analysis and decision-makers requiring accessible summaries.

## 5. Conclusions and future directions

We conducted a survey with PAM experts to highlight current challenges faced in PAM workflows, along with current strategies and emerging solutions. While experts identified nine pain points, a recurring message was that solutions already exist, in part, for most of the challenges. The problem is less about a lack of innovation than about fragmentation, duplication, interoperability issues, and uneven access. Reinventing the wheel remains common, but this approach is ultimately counterproductive. Progress will depend on building bridges among existing tools, datasets, and communities, rather than expecting that a single lab or project can develop the definitive solution in isolation.

A central theme across all categories is the need for interoperability. Whether in AI model development, metadata standards, or visualisation platforms, respondents stressed that consistent formats and modular design would allow tools to be combined flexibly, reducing inefficiency and enabling broader collaboration. This approach requires not only technical solutions but also governance structures (international standards bodies, consortia, or community-led working groups) that can coordinate efforts and ensure adoption across taxa, geographies, and user communities. Lessons from other fields, such as genomics and remote sensing, demonstrate that such coordination is possible and can dramatically accelerate synthesis.

Equity also emerged as a cross-cutting priority. While well-resourced institutions in the Global North may overcome computational and licensing barriers with local infrastructure, respondents reminded us that many high-biodiversity regions remain under-resourced. Without targeted investment, free or subsidised access to tools, and tiered support models, the global PAM community risks exacerbating inequalities in biodiversity monitoring. Ensuring that open-access repositories, user-friendly interfaces, cloud infrastructure, and training opportunities are available to researchers and practitioners worldwide is therefore essential to delivering on PAM’s promise as a truly global approach to conservation.

Looking forward, the most constructive path lies in developing a shared roadmap: a modular, open, and community-driven ecosystem for PAM. This ecosystem should integrate data storage, detection, classification, validation, and reporting in a transparent and user-friendly way, while maintaining flexibility for local adaptation and innovation. Achieving this vision will require collaboration across disciplines, bringing together ecologists, computer scientists, software developers, funders, and policymakers, and dedicated investment in infrastructure and capacity-building. If the field can avoid working in isolation, embrace shared standards, and channel collective effort into interoperable solutions, PAM has the potential to mature into a cornerstone of biodiversity monitoring, delivering actionable insights at the scales needed to confront global ecological change.

**Table 1:** Summary of key pain points in passive acoustic monitoring (PAM) workflows (ordered by their importance), with corresponding short descriptions, current strategies reported by experts, and envisioned ideal future solutions.

| Pain Point | Score | Summary | Current Solutions | Ideal Future Scenario |
| --- | --- | --- | --- | --- |
| **AI Species Identification** | 105 | Current AI models can be limited in taxonomic and geographic scope; lack of annotated datasets hinders progress. | Combine AI with manual validation; train region-specific models (e.g., BirdNET transfer learning); use data augmentation; apply statistical filters; share annotated datasets and code. | Globally diverse, open-access datasets; explainable, generalisable AI models; transparent uncertainty metrics; collaborative platforms for model- and dataset-sharing. |
| **Workflow Fragmentation** | 78 | Tools are disconnected and rarely interoperable, creating inefficiencies, errors, and difficulties scaling across projects or taxa. | Manually stitch tools together; use R/Python pipelines; modular platforms (Arbimon, PAMGuard, Raven, EcoSoundWeb); enforce metadata/file standards; SOPs for reproducibility. | Fully integrated, modular PAM platform; universal annotation/metadata standards; cloud-based scalable systems; one-stop, open-access acoustic monitoring hub. |
| **User-Friendly Software** | 65 | Many tools have steep learning curves, require coding skills, or are designed for specific taxa, excluding many ecologists. | GUIs like BirdNET, Raven, Arbimon, PAMGuard, Kaleidoscope, WildTrax; combine with R/Python scripts, Shiny Apps, workshops; collaborate with computer scientists. | Intuitive, GUI-based open-access platforms covering full workflows; modular, cross-taxon, cloud dashboards; explainable AI built into user interfaces. |
| **Manual Validation** | 42 | Checking model outputs manually is time-consuming, error-prone, and a major bottleneck for large datasets. For some understudied taxa (e.g., fish), most work is still manual | Use BirdNET Review, Raven, Triton, DetEdit, Kaleidoscope; validate subsets of detections; consensus annotation; pre-validated datasets; active learning and filtering. | Semi-automated validation with AI-assisted review, confidence scoring, consensus tools; open repositories of annotated datasets; collaborative cloud platforms. |
| **Handling, Storing, and Sharing Data** | 39 | PAM generates massive datasets that are difficult to store, back up, or share; inconsistent file formats and metadata hinder reuse. | External drives, SSDs, NAS, fragmented cloud (Box, AWS, Arbimon, WildTrax); compress files; adopt metadata standards appropriate for acoustic data; FTP servers; ship drives in low-bandwidth areas. | Centralised, open-access repositories with standardised metadata; affordable, renewable-powered cloud storage; regional data nodes; free storage for conservation projects. |
| **Standardisation and Comparability** | 35 | Lack of agreed standards for metrics, recording settings, metadata, and reporting limits reproducibility and ecological synthesis. | Precise reporting of study design and settings; core metrics for comparability; use acoustic/ISO standards; workshops and benchmarking initiatives. | Internationally agreed standards (ISO-level) for recording, metrics, and reporting; calibration protocols; open-access handbooks; global consortia to drive adoption. |
| **Computational Barriers** | 24 | Running detection/classification at scale requires computing resources unavailable to many, especially in the Global South. | Use university clusters, Google Cloud, AWS, Arbimon, EcoSoundWeb; subsample/compress data; parallel R/Python workflows; fine-tune models; optimised planning of data analyses. | Scalable, cloud-ready PAM platforms with built-in optimisation, parallelisation, and equity of access; distributed computing hubs; real-time, field-ready processing. |
| **Cost and Licensing** | 15 | High licensing fees and proprietary restrictions block access, especially in resource-limited regions; open tools exist but are fragmented. | Free/open-source tools (BirdNET, Arbimon, PAMGuard, RavenLite, R/Python); discounted licenses; collaborative license sharing; GitHub workflows. | Free, open-source PAM platforms sustainably maintained; publicly funded support; tiered systems where companies pay and academics access free; modular, well-maintained open tools. |
| **Visualisation and Reporting Tools** | 6 | Limited tools exist to generate clear visualisations, summaries, or integrated reports, complicating communication with stakeholders. | Use Raven, Audacity, PAMGuard, seewave, warbleR, monitoR; custom R/Python scripts; Power BI, Shiny dashboards; Arbimon and WildTrax for partial integration. | Integrated platforms with automated spectrograms, summaries, and reports; seamless integration of environmental/camera trap data; open, modular systems akin to Arbimon + Raven + Shiny + QGIS + eBird combined. |

## Acknowledgments

MEM thanks the support of the Australian Government through an Australian Research Council grant (project ID DE220100752). CPG was funded by the Ramón y Cajal 2024 Programme (RYC2024-048830-I) of the Spanish Ministry of Science, Innovation and Universities, funded by MICIU/AEI/10.13039/501100011033 and FSE+. We acknowledge the use of generative artificial intelligence tools to correct grammatical errors and improve the clarity of the text of an earlier version of this article. XR thanks the Belgian American Educational Foundation (BAEF) and Wallonia-Brussels International (WBI) for their support. Grants and fellowships were provided to LFT by the São Paulo Research Foundation (FAPESP #2022/11096-8) and the National Council for Scientific and Technological Development (CNPq #302834/2020-6). IM thanks the support of the Spanish National Parks Autonomous Agency (OAPN) for funding through the RarAvis project (Reference; 3377/2025).

## Author contributions

**Martino E. Malerba:** Conceptualization, Data curation, Formal analysis, Investigation, Methodology, Project administration, Resources, Writing – original draft, Writing – review and editing

**Kristian Bell:** Investigation, Project administration, Writing – original draft, Writing – review and editing

**Maria M. Palacios:** Investigation, Project administration, Writing – original draft, Writing – review and editing

**Kristen M. Bellisario, Camille Desjonquères, Alba Márquez-Rodríguez, Irene Mendoza, Christoph F. J. Meyer, Vijay Ramesh, Xavier Raick, Tessa A. Rhinehart, Connor M. Wood, Morgan A. Ziegenhorn:** Investigation, Writing – original draft, Writing – review and editing Giuseppa Buscaino, Marconi Campos-Cerqueira, Marina H. L. Duarte, Amandine Gasc, Tara Hanf-Dressler, Francis Juanes, Leandro Aparecido do Nascimento, Rodney A. Rountree, Karolin Thomisch, Luís Felipe Toledo, Mosikidi Toka, Manuel Vieira: **Writing – review and editing**

**Cristian Pérez Granados:** Conceptualization, Data curation, Formal analysis, Investigation, Methodology, Project administration, Writing – original draft, Writing – review and editing

## Supplementary Material

### 1) Expanding BirdNet models to non-avian species through transfer learning

<u>Author: Kristian Bell</u>

Centre for Nature Positive Solutions, Department of Biology, School of Science, RMIT University, Melbourne, VIC 3000, Australia

While global automated classifiers are increasingly available for birds, such tools often remain insufficient for practitioners seeking to detect non-avian taxa, rare species, or other sound types not yet covered by existing models. One practical and efficient approach to developing classifiers for new sound categories is transfer learning, which adapts a pretrained neural network to a new task by retraining it with a smaller, targeted dataset (Kath et al, 2024).

Transfer learning builds on a convolutional neural network (CNN) that has already learned to detect general acoustic features from large, diverse datasets and then fine-tunes this knowledge using recordings from the focal taxa. In many bioacoustic applications, the underlying CNN is pretrained on millions of labelled recordings from major repositories, enabling it to capture fundamental sound structures such as harmonics, frequency sweeps, and temporal patterns. For example, BirdNET (Kahl et al. 2021) was trained on millions of labelled bird recordings from repositories such as eBird, the Macaulay Library, and Xeno-canto. During fine-tuning, the model retains this capacity to represent general acoustic features while learning to associate them with the vocalisations of new species. This approach substantially reduces training time and data requirements compared with training a model from scratch, although a moderate number of labelled examples remain essential for robust performance. If the pretrained model was developed primarily for a particular taxonomic group or acoustic domain, optimal performance for other taxa may require adjustment of parameters and training settings to better match the acoustic characteristics of the target species.

This guide builds on Bell at al. (2026) and demonstrates how an accessible deep learning framework can be repurposed to train classifiers for non-avian taxa using relatively small labelled datasets. We use the BirdNET platform to illustrate this workflow as it provides a straightforward, user-friendly graphical user interface that lowers the technical barrier to entry. Many other pretrained models, including PANNs (Kong et al, 2020), YAMNet and Google Perch (van Merriënboer et al. 2025), can also be adapted in a similar transfer-learning framework, although they typically require a more code-based workflow.

As a demonstration, we show how to train a multi-species frog classifier using 28 labelled three-second call examples for each of 16 frog species (448 calls in total). The same workflow can be applied to detect other biologically or anthropogenically relevant sounds across a wide range of ecological monitoring contexts, including applications well beyond terrestrial fauna; for example, similar transfer-learning approaches have been successfully used to detect cetacean whistles (Márquez-Rodríguez et al., 2026). While evaluating classifier performance and optimising training settings can be complex and highly use-case dependent, these aspects are not covered in detail here; in most cases, the default BirdNET settings provide a robust and practical starting point for developing functional classifiers.

#### Prepare taxon-specific data

1. Assemble a labelled training set of calls for the target group. In our case, we compiled recordings of 16 frog species native to Victoria, Australia, from a combination of open-source repositories, including Xeno-canto (Xeno-Canto Foundation, 2024), FrogID (Queensland Museum, 2025), and Frog Calls of Melbourne (McNabb, 2009).
2. Recordings for each species (known as a class) should be in their own folder with clear, standardised names (**Error! Reference source not found.**). Classifier performance depends strongly on the quality and balance of training data, with larger datasets generally improving performance, but roughly equal training volumes across species recommended to minimise bias unless unequal representation is intentionally used to reflect real-world abundance (see Info Box 1.1).

**Figure G 1.1:**
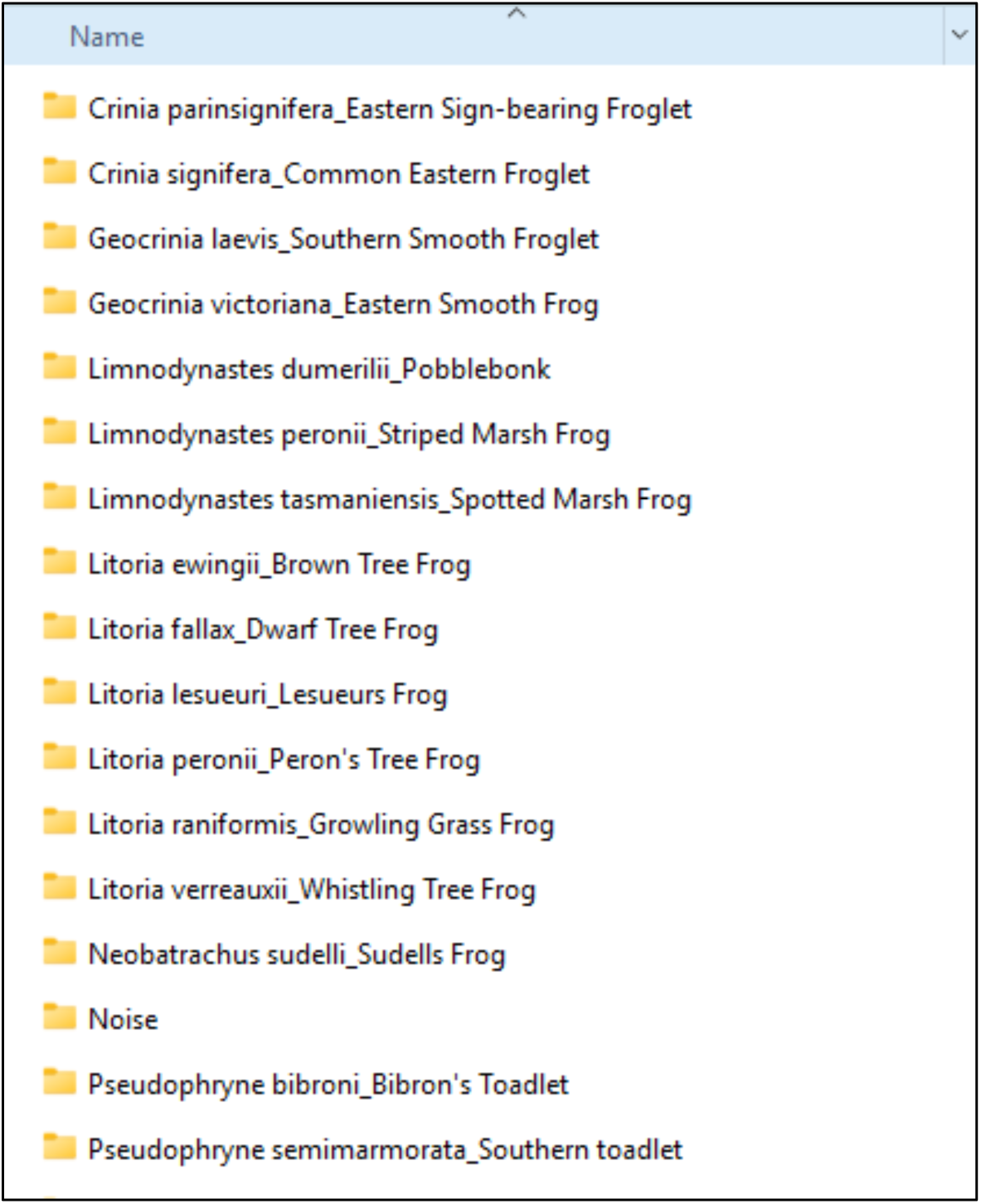
Example folder structure from Bell and Malerba (2026) for compiling training data. Each folder contains an equal number (28) of clean, 3-second audio clips for a given species or sound type. A separate ‘Noise’ folder is included to store recordings of non-target sounds (e.g., other animals, wind, rain, vehicles) used to help the model distinguish background noise from calls of interest.
3. Include representative background noise and non-target sounds to improve generalisation and reduce false positives (Info Box 1.2). In Bell et al. (2026), we included 700 example sounds of conspecific animals such as sulphur-crested cockatoos and stridulating insects, as well as other non-target noise likely to be detected given our research environment such as anthropophony (e.g., vehicles, industrial and urban activity, or human voices) and geophony (wind, rain or storms etc). Including other non-target animals helps the model learn to distinguish the focal species from acoustically similar biological sounds commonly present in the environment. Place all non-target examples in a folder named ‘Noise’.

**Info Box 1.1: Training files – how many per species?**

The quality and quantity of training data are critical determinants of classifier performance. In terms of quantity, more training files generally lead to better model performance; for CNNs, there is rarely such a thing as too much data. However, when developing a classifier that detects multiple species, it is best to maintain a roughly equal number of training files per species to minimise bias. If one species has many more examples than another, all else being equal, the classifier will be more likely to assign ambiguous sounds to the better-represented species. In some cases, a degree of bias may be desirable. For example, if certain species are naturally more common and you want detection probabilities to reflect known real-world abundance. For most applications focused on species presence within recordings, we recommend keeping training volumes roughly even across species. If achieving perfect balance is not possible, BirdNET provides automated options such as upsampling to help mitigate class imbalance (also see Guide 3: Data augmentation in PAM workflows).

**Info Box 1.2: The importance of a non-target sound class for training**

In addition to the number of training files per species, effective classifier training depends on the balance between target species sounds and non-target (background) sounds. Including a representative non-target sound class from the recording environment is critical; without it, the model is forced to assign every detected sound to the most similar species, substantially increasing the risk of false positives.

For most PAM applications, minimising false positives is particularly important because erroneous detections can generate large numbers of false records that are time-consuming to identify and remove through manual validation. Increasing the proportion of non-target training files typically makes classifiers more conservative, resulting in fewer but more reliable detections. This reduces false positives but increases false negatives, where genuine target calls are missed, leading to lower recall. This precision–recall trade-off is generally acceptable in many PAM applications because current classifiers cannot yet reliably estimate abundance, and presence–absence information is often sufficient. Moreover, the long temporal coverage of PAM deployments increases the likelihood that true presences will eventually be detected across repeated recordings, minimizing the impact of missed detections in presence–absence analyses.

The full 700-file noise class in Bell and Malerba (2026) is available at https://zenodo.org/records/17773400.

**Info Box 1.3: Clean and noisy audio training files**

When considering training data quality, it is useful to distinguish between clean and noisy recordings. Unlike the non-target sound class used to represent background noise (see Step 3), this distinction refers to the acoustic quality of individual training files. Clean recordings contain clear, typical vocalisations of the target species with minimal background noise, such as recordings of a single individual captured in isolation using high-quality equipment under controlled or semi-controlled conditions. In contrast, noisy recordings more closely resemble typical field data, often containing calls from multiple individuals or species alongside ambient sounds such as wind, rain, or anthropogenic noise. These recordings may better reflect the conditions under which classifiers are ultimately applied, as the same environmental or behavioural cues that prompt one species to vocalise frequently trigger vocalisations from sympatric species as well.

Because real-world PAM data are inherently noisy, some studies suggest that incorporating noisy recordings into training, particularly those collected using the same region, recorder type, or deployment configuration as the intended application, can improve model performance (e.g. Sasek et al. 2024). The relative emphasis placed on clean versus noisy training data should therefore be guided by the intended use case. Applications that prioritise high precision, where accurate detections are favoured over maximising recall, may benefit from weighting training data more heavily toward clean examples. Consistent with this, our own experience suggests that training on clean recordings can yield better-performing multi-species frog classifiers, even when models are subsequently applied to noisy field recordings. However, taxa that are consistently recorded in acoustically complex environments, such as fish, may derive less benefit from this approach.

#### Download and load the BirdNET GUI

4. Download the BirdNET GUI from the release page:
5. https://github.com/birdnet-team/BirdNET-Analyzer/releases/tag/v1.5.1
6. Follow the instructions to install the program once it has downloaded.

#### Create the classifier

7. After installation, open the BirdNET GUI and navigate to the ‘Train’ tab at the top of the window (Figure G 1.2).

**Figure G 1.2:**
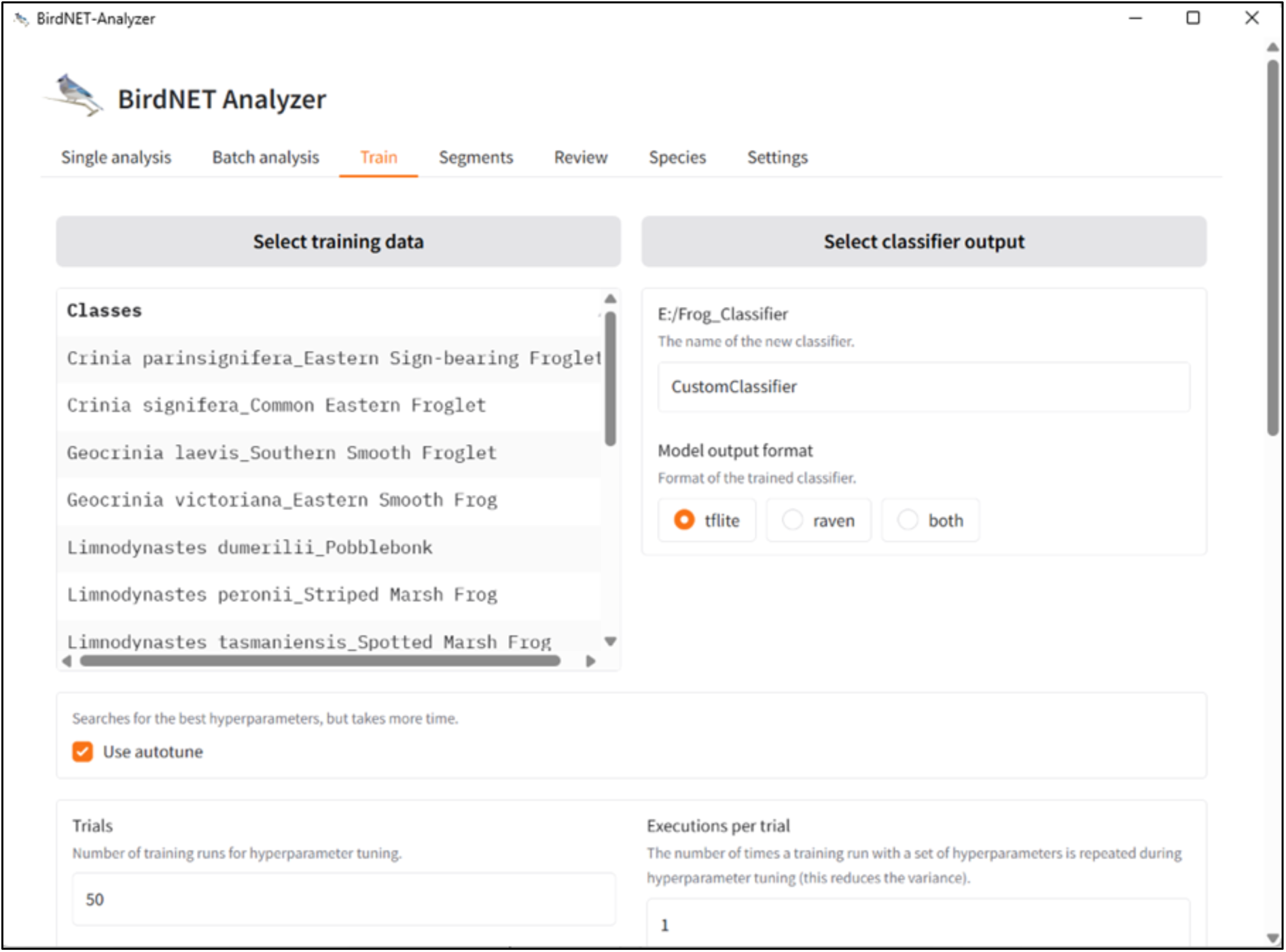
The ‘Train’ tab in the BirdNET GUI showing the setup for model training. The parent folder containing all example target sounds is selected as the training data source, and an empty, clearly named folder is designated as the output destination. The Autotune option is also enabled.

8. Click ‘Select training data’ and navigate to the folder on your computer containing all the sound files with all subfolders of example species calls (see Step 2).
9. Click ‘Select classifier output’ and create a new folder with a simple, clear name such as ‘Frog_Classifier’. This is where the resulting classifier will be saved.
10. Check the ‘autotune’ checkbox and navigate to the bottom of the window and select ‘Start training’. The ‘autotune’ option in the BirdNET training interface automatically adjusts key training parameters (such as learning rate and epoch settings) to optimise model performance for your dataset; it can help simplify training on moderately sized, balanced training sets, but may be less desirable for expert users with deep knowledge of their data who want to manually control hyperparameters or when training very large or unbalanced datasets where automated tuning might overfit or underexplore specific classes. If your training dataset is unbalanced across species, tick the ‘Upsampling mode’ option to have BirdNET resample underrepresented classes during training, helping to balance the number of examples per species. For a detailed explanation of the other training options, refer to https://birdnet-team.github.io/BirdNET-Analyzer/implementation-details/training-hyperparameters.html

BirdNET automatically reserves a portion of the training data for validation, allowing model performance to be assessed using the AUROC and AUPRC curves it provides (Figure G 1.3). In the training plots, increasing AUROC and AUPRC values indicate improving performance, while plateaus suggest limited benefit from additional training and, in some cases, the onset of overfitting. The Area Under the Receiver Operating Characteristic curve (AUROC) summarises how well the model separates positive (target species) from negative (non-target) cases across all decision thresholds, with values near 1 indicating strong discrimination and values near 0.5 indicating random performance. While AUROC is useful for comparing overall model performance, it can be insensitive to class imbalance.

The Area Under the Precision–Recall Curve (AUPRC) focuses on the positive class and captures the trade-off between precision (the proportion of detections that are correct) and recall (the proportion of true vocalisations detected). An AUPRC value of 1 indicates perfect performance, where all detections are correct and all true vocalisations are detected, whereas a value close to 0 indicates very poor performance, with detections dominated by false positives or missed true events. Because PAM datasets are typically dominated by background sounds, AUPRC is often more informative than AUROC for assessing practical detection reliability.

**Figure G 1.3:**
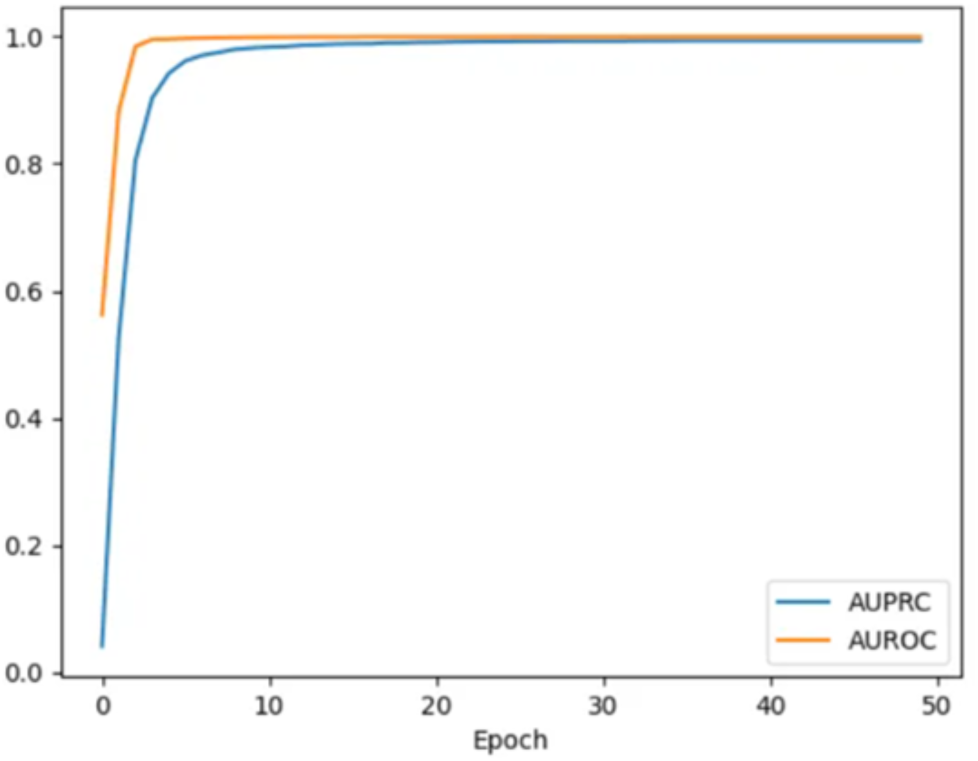
Example AUROC and AUPRC learning curves produced during BirdNET model training. Curves show validation performance across training epochs, with AUROC reflecting overall class separability and AUPRC emphasising detection performance for the target class in imbalanced datasets. Early convergence or plateauing indicates that further training provides limited performance gains. Selecting the epoch at which AUPRC first stabilises (here, around 10 epochs), rather than training to the maximum number of epochs, provides a practical balance between performance and overfitting risk.

#### Run your new classifier on other recordings

11. Go to https://www.sciencedirect.com/science/article/pii/S1574954126000506#s0080 Appendix A: Supplementary data for step-by-step instructions on how to run your newly created classifier on new data.

**Info Box 1.4: Bandpass Frequency Limits**

BirdNET allows users to ignore sounds in certain frequencies. Given that some taxa (e.g. frogs and bats) often occupy different frequency bands from birds, upon which the BirdNET CNN is based, it can be prudent to adjust the minimum and maximum bandpass frequency sliders to suit your target taxa. In our frog example, we know that our target species do not call at frequencies below 1 kHz or above 10 kHz, so we can use these values to limit the classifier analysis, and hopefully improve performance for our use case.

Congratulations - you have created your first acoustic classifier. A final, but important, step is to assess the accuracy of the positive detections produced by the classifier when applied to field recordings (a process known as validation).

The BirdNET interface allows users to extract all detected sound clips and organise them into species-specific subfolders. To do this, navigate to the ‘Segments’ tab. The programme requires three user inputs: (1) the original audio directory containing the recordings analysed by the classifier, (2) the results directory containing the output files generated during classification, and (3) a destination directory where the extracted sound clips will be saved. Create a new destination folder and give it a meaningful name; BirdNET will automatically create subfolders for each detected species.

Once these inputs are set, select ‘Extract segments’ and the programme will generate the sound clips. You can then use the ‘Review’ tab to listen to each clip and mark detections as correct or incorrect. This step is essential for evaluating classifier performance and is also useful for refining models and generating additional training data.

## 2) Building acoustic classifiers with little or no labelled training data

Author: Morgan A. Ziegenhorn

Manomet Conservation Sciences, Inc., Manomet, MA 02360, USA

Machine learning models are becoming prevalent tools for processing large amounts of PAM data, detecting and classifying vocal species at significantly faster speeds than would ever be possible with manual data labelling. However, global models like Perch and BirdNET may not perform well enough for a given PAM task without modification. Using transfer learning with these models as a base (see Guide 1), can provide a broad solution for PAM researchers by allowing researchers to leverage pretrained models for improved performance over models trained from scratch, while also potentially saving computational power (e.g., Ghani et al. 2023, Ziegenhorn et al. 2025, Márquez-Rodríguez et al. 2025).

However, such transfer learning-based models require robust training data from the target species or region. In some cases, a suitable training dataset can be developed from existing examples in public databases like Xeno-Canto (https://xeno-canto.org/), the Macaulay Library (https://www.macaulaylibrary.org/), and FishSounds (https://fishsounds.net/), but this requires that species or sounds of interest are known and that viable examples are available. In many acoustic monitoring regimes, however, researchers may be interested in targeting all soniferous species in the study area, without prior knowledge of what those species might be. Conversely, monitoring efforts may be focused on target species for which limited (or no) training data is currently available (e.g., Arctic breeding shorebirds and many aquatic and marine habitats). In this guide, we provide examples and guidance on how to use your own unlabeled PAM data to develop a comprehensive training dataset when other training data is not available.

### Step 1: Running a global model and extracting embeddings

While global models may not classify your data particularly well, embeddings produced by these models can be leveraged for signal discovery in several ways. Embeddings are a low-dimensional representation of the data learned by a given model. In 2D space, the distance between embeddings from each data segment your model classifies (e.g., 3-second segments for BirdNET, 5-second segments for Perch) should be shorter for similar segments (e.g., two songs from a singing bird) and larger for dissimilar segments (e.g., a barking dog and a singing bird). In practice, you can run your unlabeled PAM data through an existing model and extract these embeddings for further use. The exact method of extracting these embeddings depends on the global model used (see https://github.com/MZiegenhorn/birdnet-discovery and https://github.com/MZiegenhorn/birdnet-discovery and https://github.com/kitzeslab/opensoundscape for examples).

When running the global model, a brief manual inspection of a subset of detected segments using standard audio software (e.g., Raven or Audacity) is strongly recommended to confirm that biological sounds are being captured (Fig. 1). Understanding your data and the performance of your chosen model will allow you to determine whether embeddings can be successfully used for the downstream processes described in this guide. If your model is not detecting acoustic signals in your data (regardless of if classifications of those signals are accurate) then additional steps (e.g., weakly labelling a subset of data and training an intermediary classifier, see Ziegenhorn et al. 2025 for details) may be necessary.

**Figure G 2.1:**
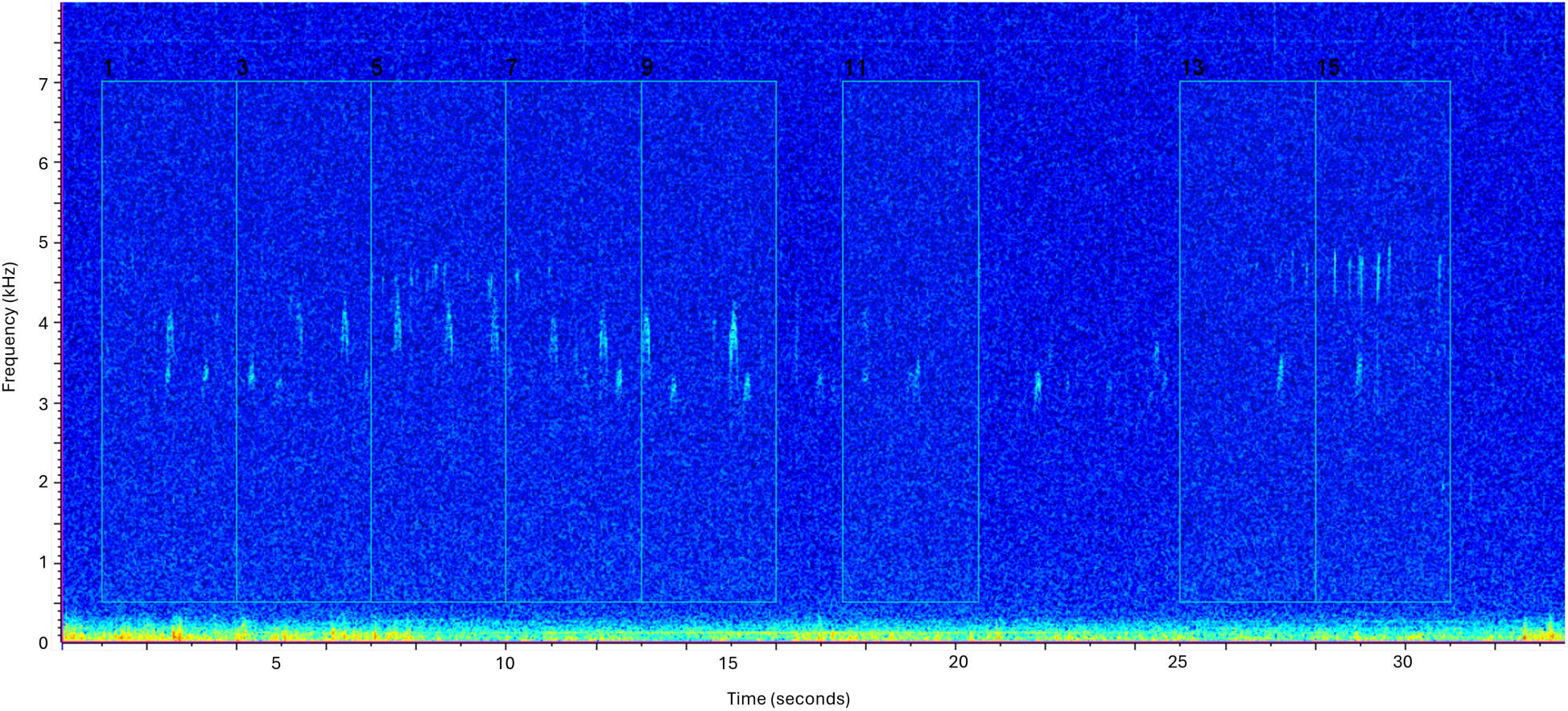
Example RavenLite spectrogram showing BirdNET detections of biological sounds, generically labelled as “bird” (white bounding boxes on the spectrogram).

Once extracted, model embeddings can be leveraged in several ways. If the target acoustic signals are known for your application, and you have access to even one good example of a given target signal, you may be able to detect the signal in your dataset without additional classification steps (Allen-Ankins et al. 2025). As outlined above, the distance between a model’s embedding of your target signal and all other embeddings of that same signal are theoretically closer together in 2D space than the embeddings of non-target sounds. Hence, embeddings with small distances from your target sound are likely detections of the target sound, which can be verified with manual review. The efficacy of this process will understandably vary based on the similarity of target and non-target sounds. Additionally, Allen-Ankins et al. (2025) noted that their method’s relatively low recall might limit its suitability for abundance estimation applications. However, this process can offer significant advantages over other signal discovery approaches in terms of efficiency, when applicable.

### Step 2: Clustering model embeddings

While useful, Allen-Ankin et al.’s (2025) approach requires researchers to know, a priori, the target acoustic signals they are hoping to detect and/or classify. If this is not known, clustering algorithms can provide a solution for signal discovery by grouping similar embeddings into clusters. Visualizations of these clusters can then be reviewed manually and given species or species group labels, allowing you to derive a comprehensive training dataset from your own PAM data (Ziegenhorn et al. 2025). One particularly useful clustering algorithm is Hierarchical Density-Based Spatial Clustering of Applications with Noise (HDBSCAN, McInnes et al. 2017). This method has recently become popular because of its performance on a variety of data types, including PAM data. Unlike other popular clustering algorithms, HDBSCAN is non-deterministic— researchers do not need to know how many clusters they expect their data to have. Additionally, HDBSCAN can readily handle uneven clusters, which is useful for PAM where cluster size may vary based on the variable commonality of different acoustic signals. Finally, this clustering algorithm can recognize noise, such that not all input embeddings are given a cluster label. This cuts down on the manual review required post-clustering to label outputs.

HDBSCAN is easy to use within the Python *scikit-learn* library, and clustering hyperparameters can be readily adjusted to fine-tune results. For example, minimum cluster size can be set very low, which may highlight rarer signals but also potentially result in a high number of similar clusters, increasing post-processing review time. You may also want to consider the pros and cons of input size—clustering one or a few short (5 minute) audio files at once is computationally quick, but results in a very large number of clusters that must be manually reviewed. Conversely, clustering many audio files (or much longer files) will result in a higher-level synthesis of clusters, but may require more RAM than is feasible with your computing set up. One option to mitigate the challenge of computing clusters across a large dataset may be to subsample your data. Determining how to subset your data, if subsetting is necessary, will be dependent on your study system and survey regime. For example, if certain times of day are more likely to contain vocalizations of birds, you may choose to focus clustering on files from those times of day (see Ziegenhorn et al. 2025 for an example of a dataset requiring subsampling). Conversely, randomly choosing files across all hours of day or survey locations may be a viable way to develop a training dataset that is representative of your study system. In addition, if the global model used to produce your embeddings includes data labels that can be trusted at a broad level (e.g., signal and noise), non-signal embeddings can be removed prior to clustering which may drastically decrease the amount of data to be clustered. An example of another subsetting method is to not include data from every study plot/acoustic instrument.

### Step 3: Cluster review and training data refinement

Clusters found using HDBSCQAN can be visualised as spectrogram images or short audio clips and reviewed manually with minimal coding in Python (Fig. 2). Example Python scripts for running HDSCAN and relevant post-processing steps are available on GitHub at: https://github.com/MZiegenhorn/birdnet-discovery. The included scripts produce spectrogram images of the data segments included in each cluster, which can be reviewed using any image-viewing software (e.g., Preview on macOS). During visual review of clusters, a spreadsheet of cluster labels can be created and updated to determine which clusters to keep, discard, or review additionally using Raven Lite or other audio-viewing software. As an example, audio clips that merit further review can be assigned to a specific ‘review’ category and then examined in greater detail using Raven Lite software (or the software of your choice) to determine their final label. This is advantageous especially when target signals may not be recognizable on sight from spectrogram images. When starting out, most clusters (that are not messy, faint, overly noisy, or of non-target signals) may be assigned to the ‘review’ class. As you progress through your cluster review, and begin defining and re-defining types, you will likely recognize some types by sight and no longer need to specifically review those files.

**Figure G 2.2:**
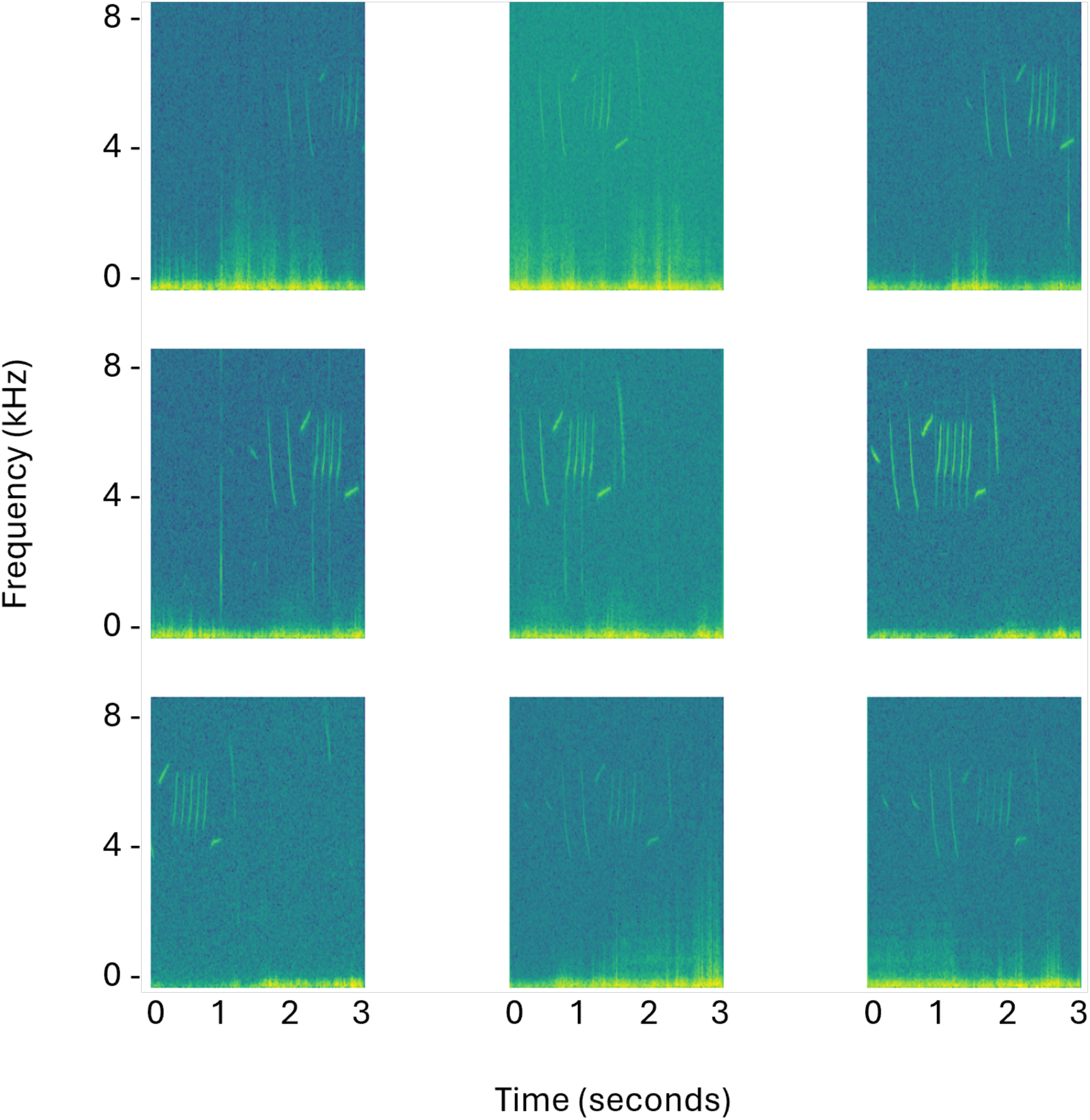
Example spectrogram images from a single HDBSCAN cluster. A PAM practitioner might conclude that this cluster is useful, as all data segments seem to contain the same signal (in this case, song from an unknown species of passerine). Further inspection of these data segments may allow for species-level identification of this signal type.

If study species in your system are not well-known to you upon starting out, you might begin with something like this:

| Deployment | Cluster # | Group |
| --- | --- | --- |
| TLSA_2023_AL54 | 0 | review |
| TLSA_2023_AL54 | 15 | LALO |
| TLSA_2023_AL54 | 16 | review |
| TLSA_2023_AL54 | 20 | review |
| TLSA_2023_AL54 | 30 | type1 |
| TLSA_2023_AL54 | 31 | type1 |
| TLSA_2023_AL54 | 32 | type1 |
| TLSA_2023_AL54 | 47 | type2 |

Here, we have one cluster that we know is Lapland Longspur (‘LALO’), several others that we would like to review (‘review’), and four clusters that we can distinguish but do not know the identity of (‘type1’, ‘type2’). As we progress through manual review of clusters in this folder and others, we can re-contextualize these clusters and assign more of them to particular species or species’ groups:

| Folder | Cluster # | Label |
| --- | --- | --- |
| TLSA_2023_AL54 | 0 | RTLO |
| TLSA_2023_AL54 | 15 | LALO |
| TLSA_2023_AL54 | 16 | PAJA |
| TLSA_2023_AL54 | 30 | WIPT |
| TLSA_2023_AL54 | 31 | WIPT |
| TLSA_2023_AL54 | 32 | WIPT |
| TLSA_2023_AL54 | 47 | AMGP |

In this case, we learned upon further data exploration that type 1 represented calls from the Willow Ptarmigan (WIPT), type 2 represented calls from the American Golden-plover (AMGP), and clusters 0 and 16 were attributed to additional species (Red-throated Loon and Parasitic Jaeger). Cluster 20 was removed after review as it was determined it was a poorer example of a species for which we already had better examples from other clusters.

The data segments distilled and labelled from this process can then be used for training a classification model that is optimized for your dataset. In Ziegenhorn et al. (2025), this process was used to create the training data for ArcticSoundsNET, which was built as a custom classifier using BirdNET as a base model. However, the training data produced by this pipeline is broadly usable for transfer learning using other base models, and, by conserving raw audio for training, is agnostic of model architecture (i.e., your model architecture need not be a convolutional neural network or utilize spectrogram images as the primary means of classification).

## 3) Data augmentation in PAM workflows

Authors: Camille Desjonqueres^1^, Alba Marquez-Rodriguez^2^, Morgan Ziegenhorn^3^ -

^1^Universite Grenoble Alpes, Universite Savoie Mont Blanc, CNRS, LECA, F-38000 Grenoble, France

^2^University of Cadiz, Institute of Marine Research (INMAR), International Campus of Excellence in Marine Science (CEIMAR), 11519 Puerto Real, Cadiz, Spain

^3^Manomet Conservation Sciences, Inc., Manomet, MA 02360, USA

Data augmentation is a generalisation strategy in machine learning in which new training samples are created by applying controlled transformations to existing data. These transformations introduce variability that reflects plausible changes in real-world conditions while preserving the original labels (Bengio et al., 2017; Krizhevsky et al., 2017). The primary objective of data augmentation is to reduce overfitting and improve model robustness by exposing models to multiple variants of the same underlying signal. This approach is particularly effective for small, unbalanced, or acoustically homogeneous training datasets. In practice, augmentation methods are most useful when computationally lightweight, so that they do not substantially increase training time.

Before applying signal-level augmentation, many PAM workflows address class imbalance using dataset-level resampling strategies. These approaches increase the representation of underrepresented classes by duplicating existing annotated samples or reducing the number of samples from dominant classes. Although resampling does not introduce new acoustic variability, it can substantially reduce bias toward common species or sound types when training data are scarce or highly imbalanced (Chawla et al., 2002; He & Garcia, 2009). Oversampling can increase the risk of overfitting, particularly when rare classes are represented by very few recordings, but it remains a simple and effective baseline when combined with regularisation or subsequent augmentation. Hence, dataset-level resampling and signal-level data augmentation address complementary aspects of model training: class balance and acoustic variability, respectively.

Several signal-level augmentation methods have proven effective in bioacoustics while generally preserving the semantic content of the original recordings (Abayomi-Alli et al., 2017). Noise addition involves adding background noise to original recordings to reduce signal-to-noise ratio or simulate different recording environments. This helps models generalise across varying acoustic conditions. Time shifting is a particularly simple and effective technique because most neural networks operate on fixed-length input segments. It varies the temporal position of labelled signals within an analysis window, encouraging models to recognise calls regardless of their exact timing. Sound mixing, also referred to as *mixup*, combines two recordings to generate more complex acoustic scenes, promoting robustness to overlapping vocalisations and background sounds (Stowell, 2022).

Additional augmentation methods include frequency shifting and time or frequency warping, often referred to as “vocal tract length perturbation”. These transformations can substantially alter acoustic cues and may affect biologically meaningful properties such as call type, individual identity, or species discrimination (Stowell, 2022). As a result, they should be applied cautiously and evaluated carefully for each use case.

Some recent architectures, particularly transformer-based models, also employ spectrogram-level techniques such as patch holding, in which portions of the spectrogram are deliberately masked to encourage learning from incomplete information. While these approaches can improve robustness, they may also reduce interpretability and are less commonly used in applied PAM workflows.

In practice, data augmentation for bioacoustics is often implemented in Python-based workflows using libraries such as *SpecAugment*, *audiomentations*, or *Kapre* (Choi et al., 2017; Park et al., 2019; Jordal et al., 2025). These libraries provide modular, well-tested implementations of many augmentation techniques and can be integrated directly into model training pipelines.

For BirdNET users, several data augmentation options are available directly during model training. Parameters such as *mixup*, *upsampling_ratio*, and *upsampling_mode* can be configured via the command line interface or the BirdNET GUI to control both dataset-level resampling and signal-level augmentation. Mixup generates additional training examples by blending pairs of recordings and their labels, helping the model generalise to more variable acoustic conditions. Upsampling_ratio controls how strongly underrepresented classes are replicated during training to reduce class imbalance. Upsampling_mode determines how this replication occurs, for example by randomly repeating existing samples or by creating augmented variants. In addition, the autotune option enables multiple training runs with different hyperparameter combinations, helping users identify augmentation settings that optimise model performance. Detailed documentation and examples are available at: https://birdnet-team.github.io/BirdNET-Analyzer/usage/cli.html#cli-docs

## 4) Coding an integrated multi-tool workflow in R

Authors: Christoph F.J. Meyer^1^ and Martino Malerba^2^

^1^Environmental Research and Innovation Centre (ERIC), University of Salford, Salford, UK

^2^Centre for Nature Positive Solutions, RMIT University, Melbourne, Australia

PAM workflows often rely on disconnected software tools that require manual file transfers between recording, visualization, feature extraction, classification, and validation tools. This fragmented process is not only time-consuming but also more likely to introduce errors at each transfer point, hindering automation and reproducibility. In particular, this approach is poorly scalable from short-term pilot studies with a limited number of audio files to long-term monitoring programmes with thousands of hours of recordings. A more robust approach is to develop an integrated R (or Python) pipeline to automate the entire workflow from raw audio through detection, classification, and validation to final analysis.

An integrated PAM workflow follows a logical continuum from field recording through to final analysis: autonomous recording units collect audio data with associated metadata, which then flows through automated detection (finding signals of interest), classification (identifying species or call types), validation (quality control and verification), and finally analysis and pattern visualization (Figure 1). Each stage builds on the previous one, with results and metadata passed forward through the pipeline, creating a traceable path from raw recordings to ecological insights. This integrated approach ensures consistency, allowing researchers to process everything from a handful of test files to large, multi-year datasets using the same code.

The following outlines the steps involved in a fully integrated workflow for PAM analysis implemented in R. It is advisable to create separate scripts for each step of the workflow, then run them in sequence from a master script.

### Step 1: Setup and file organization

A well-organized file architecture is fundamental to reproducible PAM pipelines and should follow a standardized directory structure that separates raw data from processed outputs and analysis code. The recommended structure of folders in the project directory includes */data/raw* for original, unmodified recordings, */data/annotations* to store templates or manually-labelled recordings, */detections* for detection algorithm outputs, */classifications* for species identification results, */validated* for manually verified data, */outputs* for final figures and reports, */scripts* for all R code organized by processing stage, and */metadata* for deployment information. Embedding key metadata (recording date and time, location, recorder ID) in filenames enables traceability (see Guide 7 for more details on metadata). In this context, consistent naming of files is crucial to prevent confusion and facilitate automated downstream processing. For example, use *RecorderID_Site_YYYYMMDD_HHMMSS.wav* for raw audio files. Detection outputs should reference their source file (e.g., *RecorderID_Site_YYYYMMDD_HHMMSS_detections.csv*), and intermediate files should include processing stage identifiers (e.g., *RecorderID_Site_YYYYMMDD_HHMMSS_features.csv*). Metadata should generally be stored as .csv files and capture recording context (e.g. GPS coordinates, habitat type, deployment/retrieval dates), equipment specifications (e.g. sampling rate, microphone model), and processing parameters (e.g. detection thresholds, frequency filters).

To support robust and reproducible multi-tool workflows, the use of RStudio Projects to manage file paths and project structure consistently across analyses is highly recommended. Handling audio data in R typically requires dedicated packages, such as **tuneR**, which provides core functionality for reading and writing audio files (e.g. *readWave()*, *writeWave()*). To ensure computational reproducibility, the use of **renv** is strongly recommended to record exact package versions and manage project-specific dependencies. In parallel, version control using **Git** (see Guide 10) should be adopted to track analysis scripts, configuration files, and metadata, while excluding large audio files and intermediate outputs via a *.gitignore* file.

### Step 2: Classification

Signal detection identifies temporal segments within recordings that contain acoustically structured events of interest and forms the foundation for subsequent classification and validation. In practice, users face a trade-off between using powerful pretrained models and maintaining a fully integrated workflow within a single programming environment.

#### Path A: Pretrained models (BirdNET, Perch)

Large pretrained models such as **BirdNET** provide robust classification performance across diverse soundscapes but are not currently designed to be run natively within R, and only limited Python-based programmatic interfaces are available. For most users, the most reliable approach is therefore to run BirdNET via its graphical user interface (GUI), export classification results (e.g. timestamps, predicted labels, confidence scores), and then import these outputs into R for downstream validation, summarisation, and analysis. This approach prioritises classification performance and accessibility over full pipeline integration and is widely used in applied PAM studies.

Other large pretrained models, such as **Perch**, offer greater flexibility for programmatic integration within Python-based workflows and can be embedded more directly into end-to-end pipelines. However, integration with R remains limited, and cross-language workflows typically require explicit hand-offs between Python and R.

#### Path B: Fully integrated R workflows

If a fully integrated workflow is required to be implemented entirely within R, detection must be performed using custom or classical approaches. A common strategy is to split long recordings into shorter clips (e.g. 5 s) using **tuneR**, followed by rule-based or template-based detection. Simple amplitude- or frequency-threshold detectors can be implemented using functions such as *afilter()* and *ffilter()* from **seewave**. Alternatively, template matching can be performed using **monitoR**, where acoustic templates are created from high-quality reference calls using *makeCorTemplate()*. These templates are then slid across new recordings using *corMatch()*, with detections flagged when cross-correlation scores exceed a user-defined threshold. Detected segments can be saved for subsequent feature extraction, classification, and validation within the same R pipeline.

### Step 3: Feature extraction

Following detection, acoustic events must be represented in a form suitable for classification, validation, or exploratory analysis. The appropriate representation depends on whether detection and classification are handled by an external pretrained model or implemented entirely within R.

#### Path A: Pretrained models (BirdNET, Perch)

When classification is performed using pretrained models such as BirdNET or Perch, explicit feature extraction within R is often unnecessary. These models internally compute learned representations of each detection and output species predictions with associated confidence scores. In BirdNET-based workflows, users typically import classification tables containing timestamps, predicted labels, and confidence values, which can be used directly for downstream filtering, aggregation, and validation. Where required for signal discovery or weak supervision, embeddings generated by these models can be extracted and imported into R for exploratory analysis.

#### Path B: Fully integrated R workflows

When classification is implemented within R, acoustic features must be computed explicitly. Spectral and temporal features such as peak frequency, bandwidth, duration, and dominant frequency can be extracted using **warbleR** (e.g. *specan()*) and **seewave** (e.g. *duration()*, *dfreq()*). This step produces a feature matrix describing each detected event (typically 10–20 features per detection), which forms the basis for subsequent classification or rule-based filtering.

### Step 4: Classification

Classification assigns taxonomic or functional labels to detected acoustic events, with the implementation again depending on the chosen workflow.

#### Path A: Pretrained models (BirdNET, Perch)

When using BirdNET or Perch, classification is provided directly by the model. Downstream processing in R focuses on post-classification operations, such as filtering detections by confidence threshold, collapsing predictions across overlapping time windows, or aggregating detections to higher taxonomic or functional levels. No additional model training is required unless users wish to retrain or extend classifiers using custom workflows.

#### Path B: Fully integrated R workflows

In R-based pipelines, classification is typically implemented using supervised machine learning models trained on handcrafted acoustic features. Common approaches include random forest classifiers implemented via **caret** or **randomForest**, using manually annotated training datasets and cross-validation to assess performance. For instance, the feature matrix (frequency characteristics, temporal patterns, descriptors of spectral shape) generated from a manually annotated training dataset can be used to train a random forest classifier using *train()* in the caret package with *method = “rf”*, performing cross-validation (e.g. 10-fold) via *trControl = trainControl(method = “cv”, number = 10)*. More complex models, such as convolutional neural networks operating on spectrogram images, can be implemented using **keras** when appropriate. Classification outputs should include both predicted labels and confidence scores to support transparent validation.

### Step 5: Validation and quality control

Validation is a critical step in PAM workflows, transforming preliminary detections and classifications into verified datasets suitable for ecological inference, regardless of whether detection and classification were performed using pre-trained models or fully integrated R-based approaches.

Automated validation applies rule-based filters to remove implausible detections and obvious errors. These filters can be implemented using **dplyr** functions such as *filter()* and may be based on signal-to-noise ratio thresholds to remove detections buried in noise, frequency bounds to exclude signals outside the biologically plausible range of focal species, or confidence thresholds to discard low-confidence predictions produced by pretrained models such as BirdNET or Perch. In fully integrated R workflows, similar filters can be applied to outputs from custom classifiers to remove detections near decision boundaries or with low classification confidence.

Manual validation remains essential for quality assurance, particularly for rare species, low-confidence detections, or taxa that are poorly represented in training data. Manual review can be efficiently implemented using an interactive **Shiny** application that integrates spectrogram visualisation, audio playback, and validation logging (see also Guide 5 “Developing GUI-Based Interfaces for PAM Tools”). Such applications typically rely on **tuneR** for audio handling and **ggplot2** or **seewave** for spectrogram display, and allow reviewers to prioritise subsets of detections, for example those with low classification probability or those flagged by automated filters.

Validated detections should be saved as a distinct data product, preserving links to raw recordings, detection parameters, and validation decisions to ensure transparency and reproducibility.

### Step 6: Visualisation and reporting

Data manipulation for summarising results, and indeed throughout the entire workflow, relies heavily on the **tidyverse** collection of R packages. In particular, **dplyr** functions such as *mutate()*, *filter()*, *select()*, *summarise()*, and *group_by()* are used to aggregate validated detections across temporal, spatial, or taxonomic dimensions.

For example, validated detections can be grouped by time of day or season to investigate temporal patterns of acoustic activity, or linked with deployment metadata to assess spatial patterns of species’ habitat use and preference. Aggregated results can then be visualised using **ggplot2**, enabling flexible and reproducible generation of figures such as activity curves, heatmaps, or site-level summaries.

For fully reproducible PAM workflows, it is recommended to generate automated reports using **RMarkdown** (via the **rmarkdown** package), which combines analysis code, figures, statistical results, and descriptive text into a single HTML or PDF document. This approach ensures that results derived from either pretrained-model workflows (e.g. BirdNET or Perch) or fully integrated R-based pipelines are documented in a transparent, repeatable, and shareable manner.

**Figure 1.**
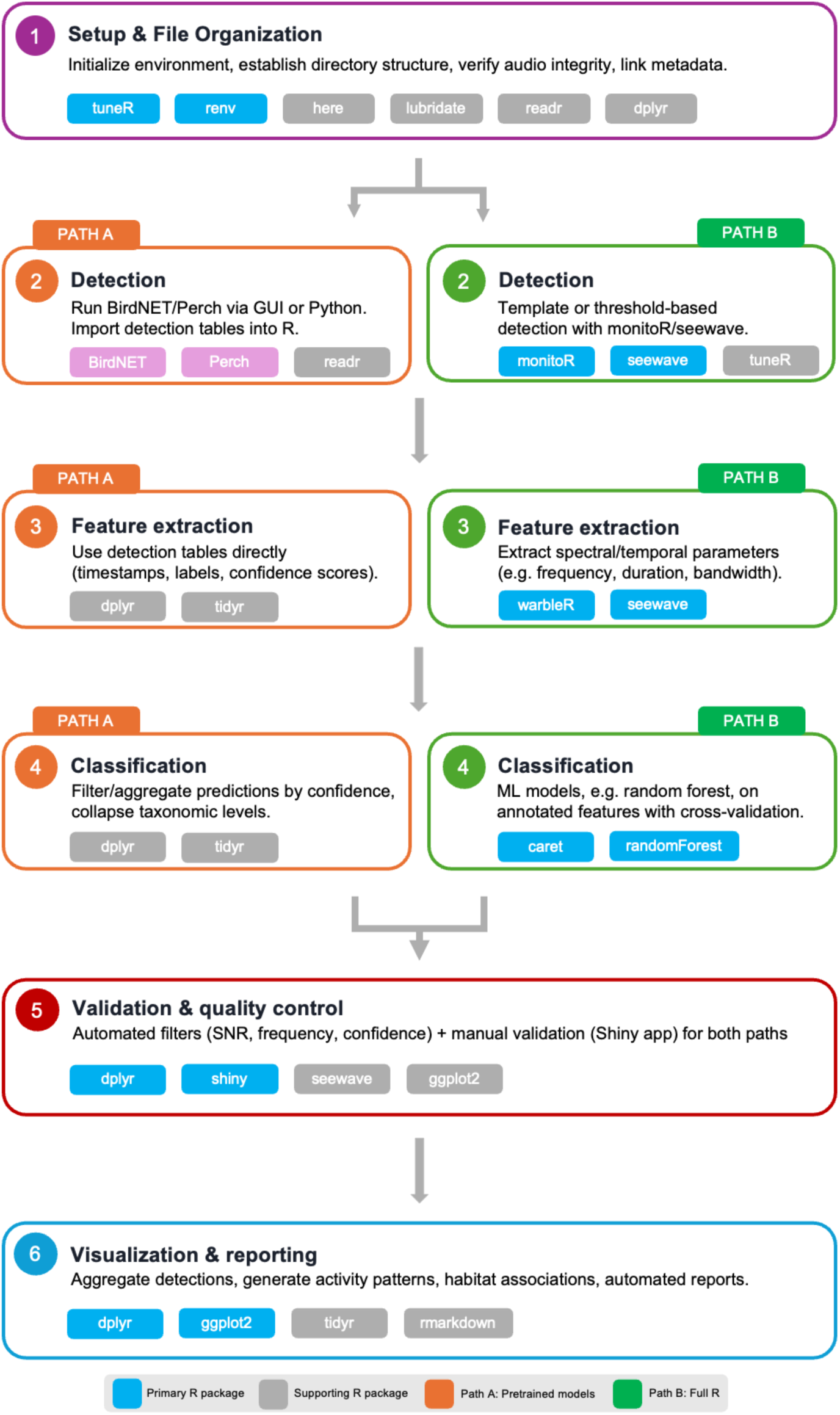
PAM workflows can follow two distinct approaches - leveraging powerful pretrained models with R used primarily for post-processing or implementing end-to-end analysis entirely within R while maintaining the same core workflow structure.

## 5) Developing GUI-based interfaces for PAM

Authors: Kristen M. Bellisario^1^ and Martino E. Malerba^2^

^1^Purdue University, West Lafayette, IN 47906, USA

^2^Centre for Nature Positive Solutions, Department of Biology, School of Science, RMIT University, Melbourne, VIC 3000, Australia

Integrated workflows for passive acoustic monitoring are often implemented through code-based pipelines, particularly in R or Python, because these environments offer flexibility, transparency, and access to a wide ecosystem of analytical tools (see Guide 4 “Integrating multi-tool workflows in R and Python”). However, code-based workflows can represent a significant barrier for users without programming expertise, including practitioners, managers, students, and citizen scientists, who engage with PAM at specific decision points rather than across full analytical pipelines.

To increase accessibility while preserving analytical rigor, there is a growing trend toward the development of graphical user interfaces (GUIs) that expose key PAM functionalities through interactive visual tools. Well-designed GUIs allow users to engage with complex workflows such as detection validation, threshold selection, annotation, and exploratory analysis without writing code, while still producing outputs that can be integrated into reproducible analytical pipelines. GUI-based tools should not be viewed as replacements for reproducible analytical pipelines, but rather as complementary interfaces that support specific tasks within them, including reviewing automated outputs, validating detections through human inspection, and exploring emerging ecological patterns.

Effective GUI-based tools for PAM typically focus on a limited number of well-defined tasks rather than attempting to replicate entire pipelines. One class of PAM GUIs focuses on post-processing, reviewing and validation of outputs generated by BirdNET. One example is **birdnetTools**, an R package that includes an interactive Shiny application to guide users through common decision points such as setting confidence thresholds and filtering detections. Core functionalities include importing BirdNET output tables, filtering detections by species, confidence, and date or time, and visualising temporal patterns of detections. The BirdNET Shiny Real-Time Frontend (“**shiny.rtf**” Weidlich-Rau et al. 2025) offers similar interactive review capabilities with an emphasis on real-time visualisation of classifier outputs. By embedding these steps in a GUI, both tools enable users to explore and validate model outputs interactively while retaining compatibility with reproducible, script-based workflows. This type of interface supports decision-making about automated detections prior to downstream ecological interpretation or statistical modeling, helping users ground-truth classifier outputs before further analysis (See: https://birdnet-team.github.io/birdnetTools/articles/birdnetTools.html).

**Figure G 5.1:**
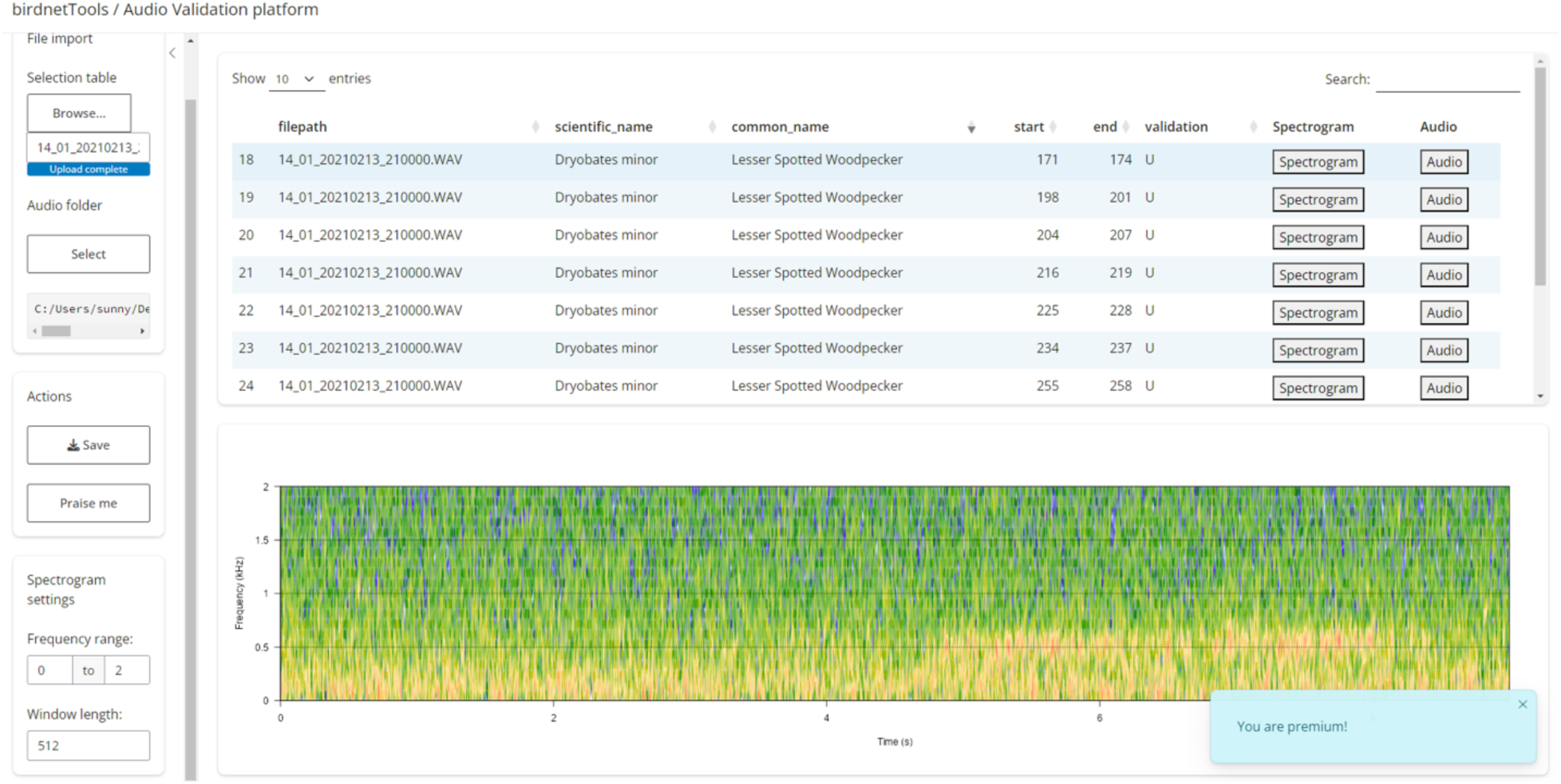
Example of the birdnetTools GUI to do post-processing of outputs from BirdNET.

Another class of PAM GUIs focuses on manual annotation and validation of short audio segments. **Audiotate** (Audio Segments Validator–Annotator) is a lightweight, Python-based application designed for efficient review and labelling of short clips, including unlabeled samples or predictions generated by deep-learning models such as BirdNET or Perch. Audiotate provides an intuitive interface that allows users to load structured folders of audio files, listen to individual segments, view spectrograms, assign or edit labels (either predefined or user-defined), add optional comments, and export annotations to a CSV file. These outputs can then be used directly for downstream analysis, training data creation, or model refinement. Comparable tools exist in other PAM workflows, such as Nature and Energy Audio Labeller (“**NEAL**” Gibbons et al. 2023), which also provides a graphical interface for listening to recordings, inspecting spectrograms, and manually assigning species labels to audio segments during dataset preparation and validation. Such interfaces explicitly reintroduce human expertise into PAM workflows, addressing uncertainty that cannot be resolved through automated filtering alone, and play a critical role in detector validation and dataset curation without requiring programming expertise (See https://github.com/SEANIMALMOVE/Audiotate_Audio-Segments-Validator-Annotator).

**Figure G 5.2:**
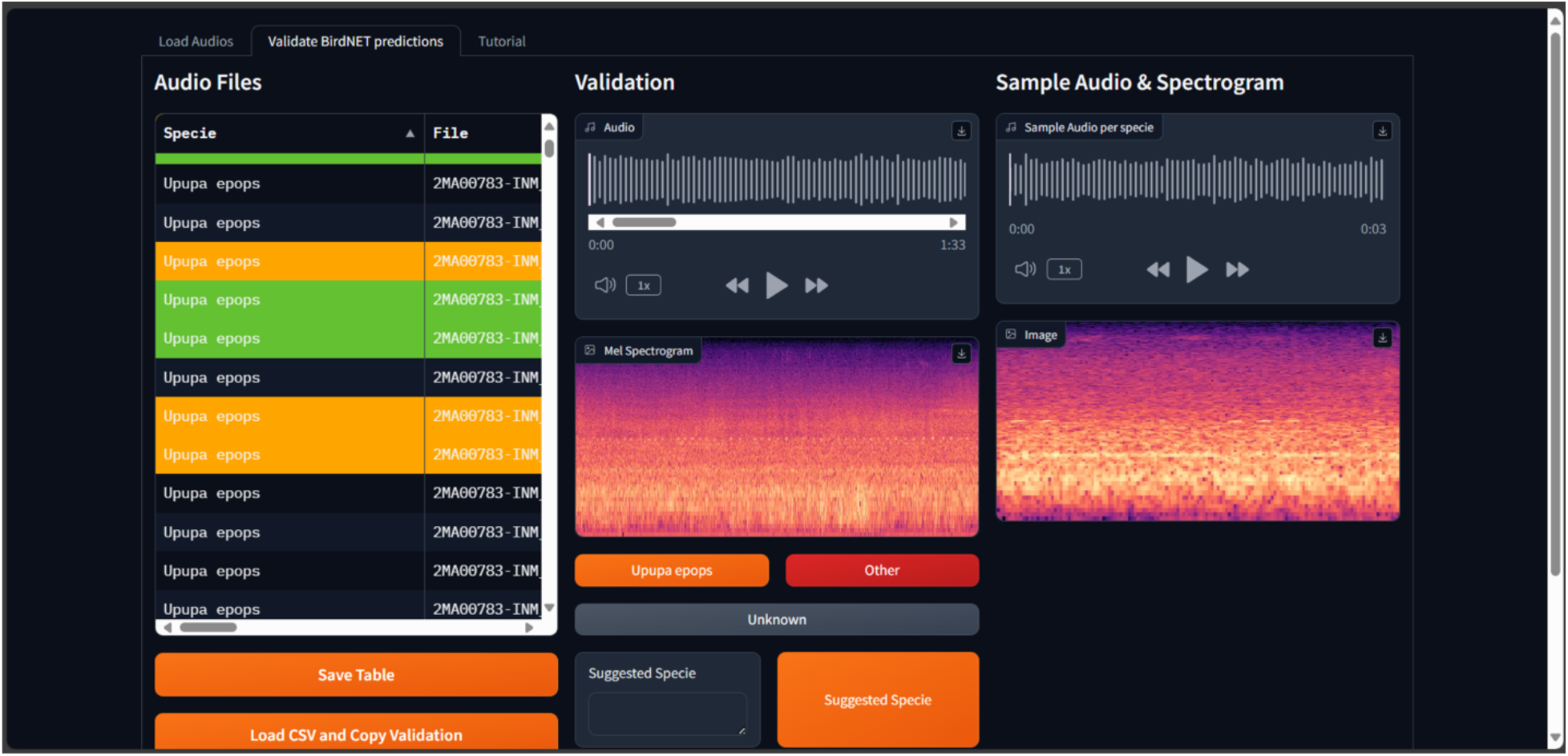
Audiotate GUI to validate audio from AI classifiers by visualizing spectrogram and listening to audio files.

A third class of GUI-based tools focuses on rapid visualisation and exploration of large PAM outputs to support ecological interpretation. For example, **ChirpCheck** (Peña Leon et al., in review) is an open-source tool for code-free exploration of BirdNET outputs, allowing users to upload classifier outputs via a drag-and-drop interface, normalise detections, and explore species composition, temporal patterns, and diel or seasonal heatmaps through preconfigured interactive dashboards. Similar exploratory interfaces are provided by **Arbimon** (Aide et al. 2013), which integrates large-scale acoustic data storage, automated detection pipelines, and interactive dashboards for exploring temporal patterns of species activity, and **soundscapeR** (Luypaert 2024), an R Shiny interface for visualising soundscape indices across long-term datasets. These tools also help expose data-quality and pipeline issues, such as missing intervals, abrupt drops in detections, or artefacts linked to recorder failure. Confidence-threshold filters and species selectors allow users to test the sensitivity of apparent patterns to classifier settings. Importantly, tools such as these are not intended to replace statistical modelling or detector validation workflows, but rather complement them by providing a rapid initial assessment and lowering the technical barrier to ecological sensemaking from large PAM datasets.

**Figure G 5.3:**
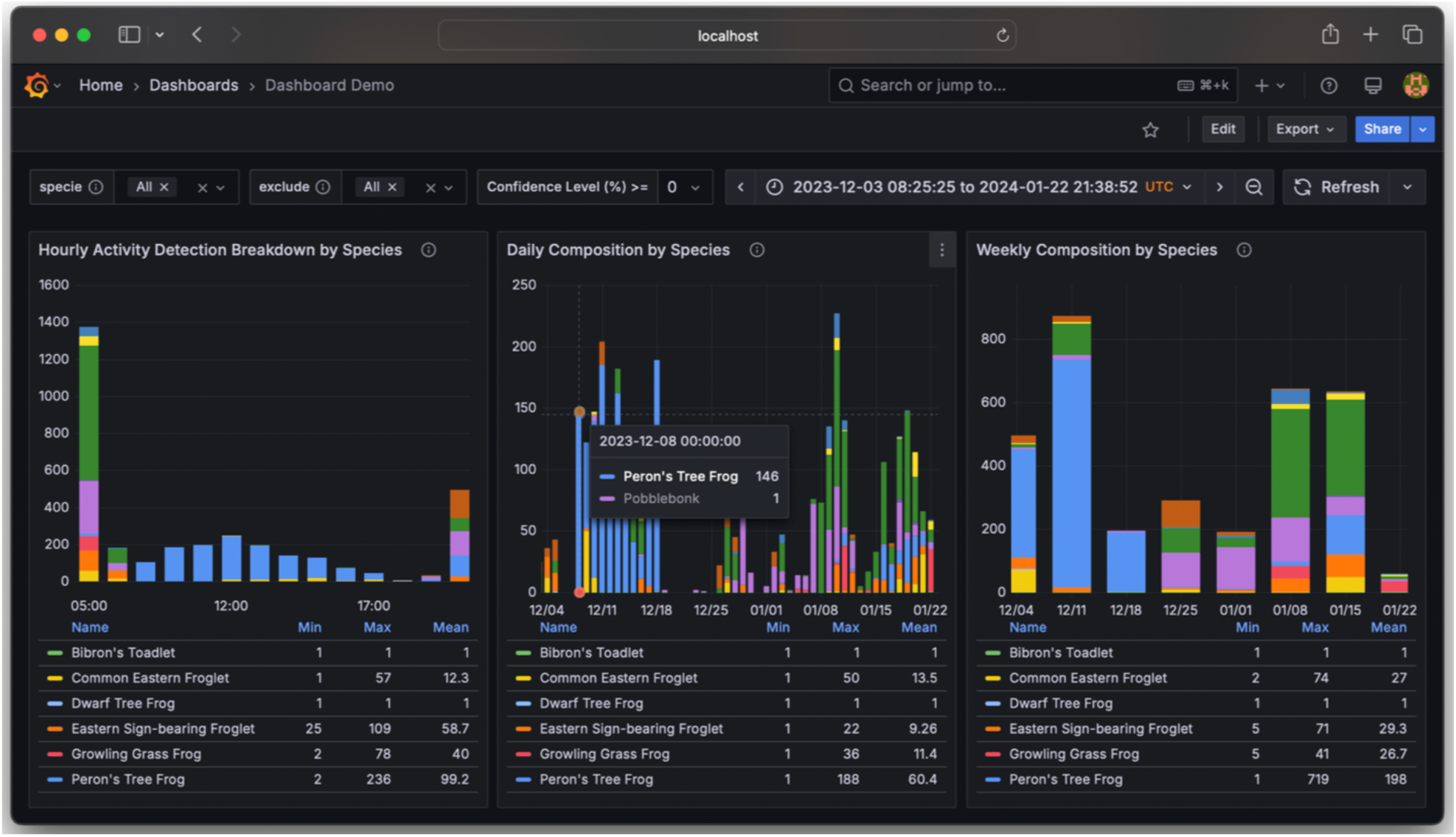
ChirpCheck dashboard panels from Garcia et al. (2025) visualising temporal patterns in species detections derived from BirdNET output files. The platform ingests multiple BirdNET-generated CSV datasets, aggregates and updates them, and displays changes in species occurrence over time through interactive, user-friendly panels designed for ecological interpretation and reporting.

Collectively, these tools form an intermediate layer between raw acoustic recordings and fully scripted analytical pipelines, supporting key analytical decisions without requiring users to write code. By focusing on discrete, well-defined tasks, they lower technical barriers while preserving transparency, traceability, and compatibility with reproducible workflows. While open-source interfaces are increasingly available, tools that support rapid, code-free ecological interpretation of terrestrial PAM data remain comparatively limited. As PAM continues to expand across research domains, well-designed open-source GUIs will play an essential role in establishing accessible baseline tools that accelerate adoption while maintaining analytical rigor.

## 6) Accounting for classifier errors to make correct ecological inferences

Authors: Connor M. Wood^1^ and Tessa A Rhinehart^2^

^1^ K. Lisa Yang Center for Conservation Bioacoustics, Cornell Lab of Ornithology, Cornell University.

^2^ Department of Biological Sciences, University of Pittsburgh.

Why should you or anyone else trust your detector’s predictions? Without expert validation, no one should. Here are several approaches:

- **Review of all predictions above a threshold.** The simplest but most time-consuming approach is to review all the predictions above a given threshold and use only the correct predictions (i.e., confirmed observations) in your subsequent analyses. In the case of legally protected species or other high-stakes applications, it may be necessary to only work with confirmed true positive observations. To some extent, you can set that minimum threshold based on how much time you can afford to allocate to manual verification. If you can review 500 predictions per hour and you can spend a full week on the task, you can set a threshold that yields 20,000 predictions.
- **Calibration.** One simple approach is to review a random selection of detector predictions spanning the entire dataset and then relate the binary outcome (correct or incorrect) to the continuous prediction score via logistic regression (Wood and Kahl 2024, Navine et al. 2024). You can then solve your regression equation for a desired probability that a detector prediction is correct. There are several important aspects to this approach.

First, you must review enough predictions that the variation in your dataset is accurately reflected, which makes the application of that outcome∼score relationship to the entire dataset appropriate. How many is “enough” will depend on your dataset and how much variation (in both the soundscape and the target signal) it entails. Furthermore, accuracy may vary between sites due to changes in the attributes of data (i.e., *domain shift,* described below), potentially introducing false positives and making inferences at particular sites invalid.

Second, the probability of a true positive (i.e., Pr(true positive)) for a single clip is not equivalent to precision within your thresholded data despite its conceptual similarity. For example, the regression may predict a clip’s Pr(tp) is 0.95 at a score of 0.90, but if most of the predicted positives get a score of 0.99, precision may be much higher than 0.95. Ultimately, after you have implemented this approach, you can filter your detector outputs to a desired level of quality (e.g., everything with a Pr(tp) < 0.99 is discarded) and treat the retained predictions as observations.

You can calculate an expected upper bound on the number of false positive predictions you have. For example, if you keep 1,000 clips after filtering your data to the threshold at which pr(tp) < 0.99, you are expected to have 0.01*1000 = 10 false positive clips. How much of a problem this is for your data depends on many factors. For example, if all the false positives are “clumped” at one site, this may not impact your ecological conclusions. If you only have 20 occupied sites and the false positives are spread across 10 other sites, then 10 false positives could significantly impact your conclusions.

Generally, such site-level false positives—where a species is predicted to be present at sites where it does not occur—are the most likely issue to add noise to, or even bias, ecological conclusions, especially those made from occupancy models. For an example of how these problems alter your results, see Berigan et al. (2019); for an overview of the many false-positive solutions developed for occupancy models see Clare et al. (2021). Thus, all efforts should be taken to avoid them, including hybrid thresholding/review approaches (see below for examples).

- **Classifier-guided review.** Depending on the scale of the dataset, your validation time could be better spent in a structured (i.e., non-random) way. A good approach is to use “classifier-guided review” to build a detection history (<u>Katsis et al. 2025</u>). Occupancy models generally require multiple visits per site in order to fit the model. The workflow goes like this:

- Subset predictions from each site into survey periods (e.g., eight week-long survey periods throughout the study season)
- Perform some preliminary listening to select a score threshold below which you will not review clips, where the chance of a true positive becomes very low
- Sort the clips scoring above this threshold from each combination of site & survey period from highest-scoring to lowest-scoring and start listening from the highest-scoring clips down.
- If you verify the sound of a species in a clip, give that site-survey period combination a detection (1) in your occupancy modeling detection history, then stop reviewing clips from that site-survey period combination. This can save a lot of review time!
- If you hit your pre-determined threshold without finding the species’ sound in a clip, give that site-survey period combination a nondetection (0) in your occupancy modeling detection history.
- If a site has no clips above the threshold for a given site-survey period combination, give that site-survey period combination a nondetection (0) in your occupancy modeling detection history.
- **Hybrid calibration & classifier-guided review.** For massive datasets, the complete site-survey period review may be impractical but a hybrid approach can still be applied by performing the logistic regression approach described above, then using classifier-guided review to remove site-level false positives within the data (see above). For example, the detector outputs could be filtered to a particular level (say, pr(true positive) ≥ 0.95) and then the highest-scoring predictions from each location are manually reviewed. Once some quantity of correct predictions is confirmed at a site, you can move on to the next site. For example, you may be satisfied with one confirmed positive clip if you just want to be sure the species is present, or 10 confirmed positive clips separated by at least 7 days if you want to remove sites where the species was transient during migration. If a site never reaches your predetermined threshold of confirmed clips, then consider the site to be unoccupied, and replace its entire detection history with nondetections (zeroes). In this way, there are no site-level false positives, even if some of the predicted positive clips at sites where occupancy has been confirmed may be incorrect. This would overall inflate the prediction of detection probability, and potentially underestimate the number of occupied sites.
- **Threshold-free approaches.** Another alternative is using “threshold-free” approaches that use a combination of the underlying distribution of machine learning scores from each site and some amount of validation data to predict values such as site occupancy (Rhinehart et al. 2022, Katsis et al. 2025) or number of calls within a region (Navine et al. 2024). These approaches have not yet been widely applied, and are subject to the same domain shift biases that the calibration approach is.

Automated detectors create new research possibilities by accelerating the search for target signals in bulk audio datasets. However, manual validation remains a critical step in the appropriate use of such tools. Without a human in the loop, a particularly pernicious type of error can be introduced: the ones you do not know about.

## 7) Using metadata standards for acoustic data management

Authors: Alba Marquez-Rodriguez^1^ and Irene Mendoza^2^

^1^University of Cadiz, Instituto Universitario de Investigacion Marina (INMAR), Campus de Excelencia Internacional del Mar (CEIMAR), Puerto Real, Cadiz, Spain

^2^ University of Sevilla, Dept. of Plant Biology and Ecology, Sevilla, Spain

Despite the rapid expansion of passive acoustic monitoring across ecosystems and taxa, the lack of standardised metadata remains a major barrier to data interoperability and reuse. While recent technological advances have made acoustic recorders more affordable and widely deployed, it is still challenging to find a standardised procedure for describing, storing and sharing resulting recordings. As a result, PAM datasets are often difficult to integrate across projects, platforms, or regions, limiting opportunities for synthesis and reducing the long-term scientific value of collected recordings.

These challenges directly relate to the FAIR data principles, which state that data should be Findable, Accessible, Interoperable, and Reusable (GO FAIR, 2025). Metadata play a central role in achieving these principles by enabling dataset discovery through standardised descriptors, clarifying access and licensing conditions, supporting interoperability through shared vocabularies (e.g. consistent and clear naming conventions), and providing the contextual information required for meaningful reuse. Without consistent metadata, even well-curated acoustic datasets risk becoming hard to access, underused, and lost over time. Hence, establishing common approaches to metadata is a foundational step toward maximizing the scientific and conservation value of PAM data.

### What are metadata and why do they matter?

Metadata, literally “data about data”, provide the contextual information needed to interpret, reuse, and integrate acoustic recordings. They describe how, when, and where recordings were collected, as well as the technical conditions under which they were made. Without this context, acoustic files are difficult to interpret correctly, and their scientific value is substantially reduced.

In PAM, metadata can be broadly divided into internal and external types. Internal metadata are embedded within the audio files themselves and typically capture technical attributes such as file format, sampling rate, bit depth, duration, and channel configuration. These metadata are essential for ensuring that recordings can be correctly processed and analysed.

External metadata are stored separately in accompanying files, for example in JSON, CSV, XML, or text formats. They provide contextual information that is not contained in file headers, including recording location, deployment details, recorder and microphone identifiers, observer or project information. When machine-learning routines are applied to, these external metadata, link to manual annotations or automated detection and classification outputs. Together, internal and external metadata form a complete description of an acoustic dataset, enabling recordings to be interpreted, shared, and reused across projects and analytical workflows.

### Existing standards and their application to acoustics

A range of metadata standards has been developed in biodiversity and ecological monitoring to support data interoperability and reuse. **Darwin Core (DwC)** is the most widely used standard in biodiversity informatics and provides a well-established framework for documenting species occurrences, sampling events, and associated observations (Wieczorek et al., 2012). DwC underpins major biodiversity infrastructures such as GBIF (GBIF, 2025), enabling large-scale data discovery and synthesis. However, DwC was not designed with bioacoustic data in mind and it lacks dedicated fields for continuous recordings, acoustic annotations, or model-derived detections, which limits its direct applicability to PAM datasets.

A practical example of how DwC can be adapted for PAM is provided by the BIRDeep project, a regional bird monitoring initiative in Doñana National Park, southern Spain. BIRDeep employs a Darwin Core–based metadata table to capture project- and site-level information, including geographic coordinates, species lists, and campaign details (Márquez-Rodríguez et al., 2025). Because DwC does not natively support acoustic annotations, each recording is accompanied by a separate file containing identification annotations, together with a comprehensive CSV file linking audio paths to annotations and species information. This hybrid approach preserves compatibility with biodiversity data infrastructures while accommodating key requirements of acoustic monitoring. At the same time, it highlights current limitations, as technical audio parameters such as sampling rate and bit depth are not yet systematically recorded.

In contrast to other ecological monitoring domains, PAM has comparatively few dedicated metadata standards. **GUANO** is a notable exception, developed specifically for bat acoustic monitoring, and defines metadata fields for recording settings, deployment information, and detection outputs embedded directly within audio files (Riggs, 2018). Another example is the **Tethys standard**, which was designed for oceanic passive acoustic monitoring and provides a comprehensive framework for managing large acoustic archives (Roch et al., 2013, 2016). However, its structure is less readily transferable to terrestrial PAM without substantial modification.

By comparison, several mature standards exist for categorised image data collected with camera traps. **Camtrap DP** (Bubnicki et al., 2023), for example, provides a robust and widely adopted data model for camera-trap studies, with standardized representations of deployments, events, media files, and species detections. Its success in large-scale biodiversity initiatives demonstrates how structured, hierarchical metadata can support interoperability across projects and long-term data reuse. Importantly, Camtrap DP was explicitly designed to extend Darwin Core concepts to sensor-based monitoring data, offering a useful conceptual and technical foundation for other modalities such as acoustics.

Building on the Camtrap DP data model, the recently proposed *Safe and Sound* initiative adapts established biodiversity metadata practices, drawing on both Darwin Core and Camtrap DP, to the specific requirements of PAM (Wiel et al., 2026). Rather than introducing a new standard, it defines acoustic-specific entities, relationships, and best practices, explicitly distinguishing between deployment-level, recording-level, and annotation-level metadata. The framework provides guidance for linking continuous audio files with derived products such as manual annotations, automated detections, embeddings, and model outputs, while maintaining interoperability across platforms and workflows. By formalising how acoustic-specific information (e.g. recording effort, sensor configuration, temporal coverage, and annotation provenance) should be structured and documented, *Safe and Sound* offers a practical pathway toward FAIR-compliant PAM datasets without requiring researchers to abandon existing biodiversity data infrastructures.

Likewise, the **COCO** model, originally developed for computer vision tasks (Lin et al., 2014), defines a flexible hierarchy linking media objects to annotations, labels and derived outputs. The **COCO-CameraTraps** extension illustrates how this structure can be adapted for ecological applications, supporting large-scale annotation, model training, and benchmarking workflows in wildlife monitoring (Beery et al., 2018). Alongside annotation-focused formats, standards such as **Cam trap DP** (Bubnicki et al., 2023) address the complementary need for interoperable data packaging and sharing across repositories - illustrating that no single schema serves all stages of the monitoring pipeline. As artificial intelligence becomes increasingly central tool to automate biodiversity monitoring, such annotation-centric data models are particularly relevant for PAM, where recordings are continuously re-analysed using evolving algorithms. Adopting metadata frameworks that explicitly link raw media, annotations, and model outputs will be critical to ensuring transparency, reproducibility, and long-term reuse of acoustic data in AI-driven ecological research.

### Essential metadata fields in PAM

While existing standards provide useful guidance, no dedicated and widely adopted metadata framework currently exists for bioacoustic datasets. This gap is becoming critical as artificial intelligence plays a growing role in PAM workflows, where recordings are repeatedly re-analysed using different models, parameters, and validation strategies. A suitable metadata framework must therefore capture not only project-level and file-level information but also annotations and AI-derived outputs in a way that supports model training, evaluation and reproducibility. Ideally, such a standard would align with Darwin Core for interoperability with existing biodiversity infrastructures yet extend its scope to accommodate the unique characteristics of acoustic data, including multi-layered annotations, automated detections, and temporal signal properties.

To ensure consistency, interoperability, and reproducibility, acoustic datasets should be organized using a hierarchical metadata structure that captures information at multiple levels (project, license, file, annotation, automated identification, and category levels; Figure 1). This approach allows projects of different scales and analytical complexity to remain compatible while accommodating diverse annotation methods and processing pipelines.

**Figure 7.1.**
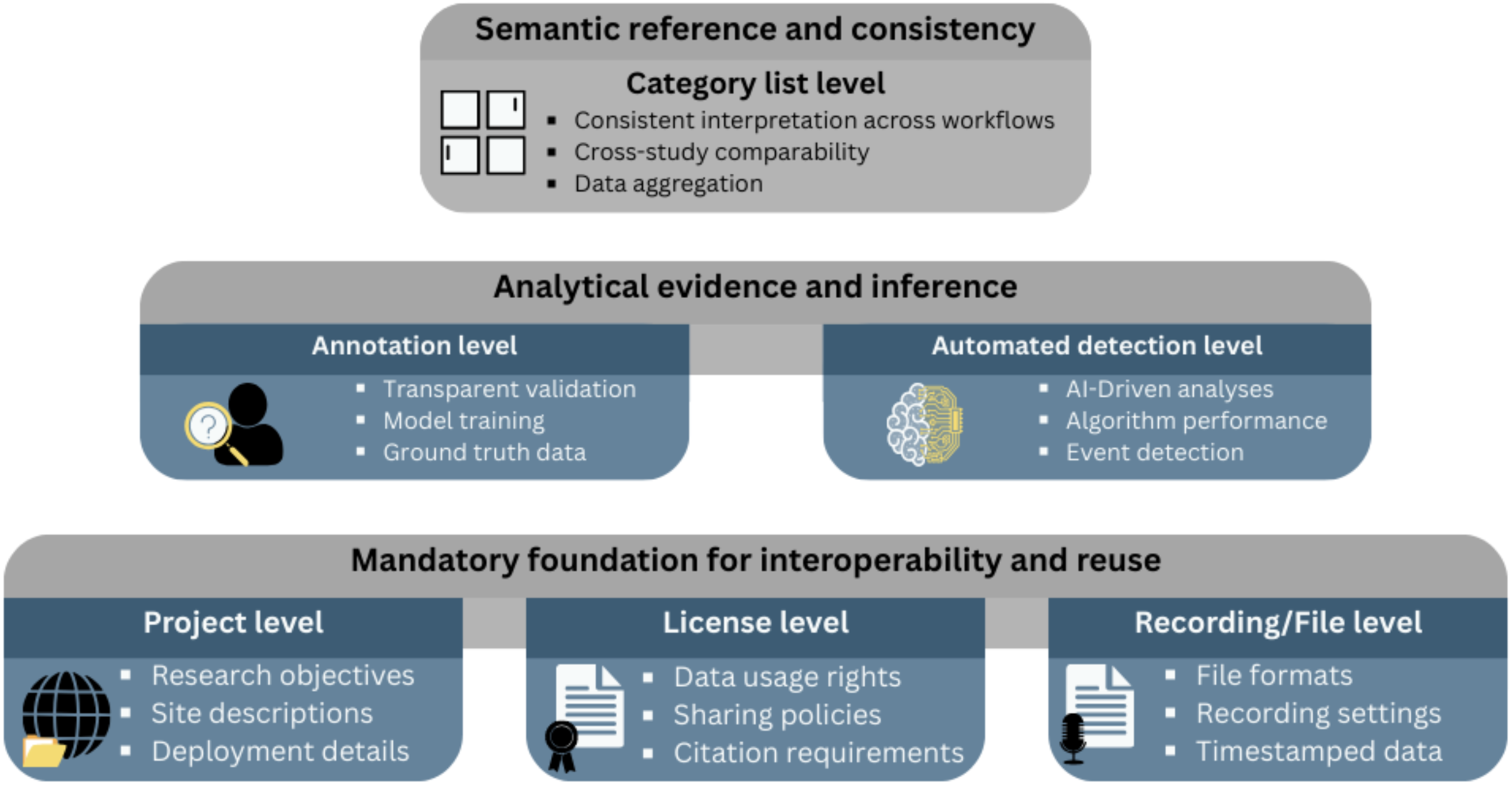
Conceptual hierarchy of essential metadata fields in PAM. The pyramid illustrates three complementary layers of metadata organization: the basis are those mandatory fields that provide foundation for interoperability and reuse (project, license, and file-level metadata); an analytical evidence and inference layer linking recordings to manual annotations and automated model outputs; and a semantic reference layer that ensures label consistency across analyses.

At the foundation of this structure there are three mandatory metadata levels that every PAM project should include. The **project level** describes the overall monitoring campaign and data organization, including the project name, objectives, geographic scope, and deployment dates. The **license level** specifies data ownership, access rights, and data-sharing conditions, ensuring compliance with open-data policies and ethical requirements. The **file level** contains core information for each recording, including date and time, geographic coordinates, locality name, recorder identifier, recording settings such as sampling rate, gain, and filter bands, and hardware specifications such as recorder and microphone models. Together, these three levels define the minimum and mandatory information required for acoustic data to be FAIR-compliant and reusable across repositories.

Additional metadata layers become relevant once annotation and analysis are performed. The **manual annotation level** captures expert-verified identifications of sounds, including start and end times, frequency range, species name, call type, annotator identity, and validation method. These annotations represent the most reliable information available and should serve as ground truth for training and evaluating automated models. The **automated or semi-automated detection level** records outputs generated by algorithms, including the source file, model name (for example BirdNET, Perch, or a custom model), model version, relevant hyperparameters, prediction start and end times and frequency range, predicted species or class, confidence score, and, where applicable, constraints such as restricted species lists and flags indicating expert validation. Finally, the **category list level** provides a summary of all species or acoustic classes represented in the dataset, based on manual annotations and or validated automated detections.

This hierarchical and modular structure, inspired by the COCO camera-trap data model, provides a flexible yet standardized approach to organizing bioacoustic metadata. By adopting such practices, PAM projects can improve data transparency, facilitate large-scale synthesis, and establish a foundation for the development of a future global metadata standard tailored specifically to acoustic biodiversity monitoring.

### Ethics and security

Ethical and security considerations are fundamental to responsible acoustic data management. Metadata can inadvertently expose sensitive information, such as the exact locations of threatened species or recordings containing human voices, posing risks to both wildlife and individuals. It is therefore essential to compliance with relevant data protection regulations, such as national Data Protection Laws or institutional ethical standards.l. To mitigate these risks, researchers should adopt good practices including the anonymization or spatial generalization of sensitive site coordinates, the automatic detection and removal of human speech from recordings, and the implementation of controlled access systems for data storage and sharing. By integrating these measures into metadata frameworks and data management workflows, PAM practitioners can ensure that open and reusable datasets remain ethically sound and secure.

### Conclusions and future perspectives

The development of a dedicated metadata standard for bioacoustics is essential to ensure the interoperability, transparency, and long-term value of PAM datasets. The previously developed standard for biodiversity data, Darwin Core, is insufficient for PAM datasets. Other existing frameworks from related domains, such as camera trapping, provide useful foundations, but they must be adapted to reflect the distinctive characteristics of acoustic monitoring, including continuous recordings, taxonomic breadth, and complex, multi-layered annotation workflows. Establishing a community-driven and openly governed standard will be critical for enabling meaningful data sharing, comparison, and synthesis across projects and ecosystems. This is still a pending task, although initiatives such as Safe and Sound project are giving very valuable solutions

Beyond supporting data management and reuse, robust metadata practices are increasingly relevant to analytical workflows themselves. Recent work suggests that incorporating contextual metadata as model inputs can improve the performance, robustness, and interpretability of artificial intelligence approaches applied to ecological data (e.g., Jeantet & Dufourq, 2023). Strengthening metadata standards therefore serves a dual purpose: facilitating open and collaborative infrastructures for PAM while also contributing to more reliable and context-aware ecological inference. As PAM continues to scale globally, investment in shared metadata standards will be a prerequisite for translating large acoustic datasets into actionable biodiversity knowledge.

## 8) Reproducible scientific workflows for PAM

Authors: Xavier Raick^1,2,3^ and Vijay Ramesh^4^

1. K. Lisa Yang Center for Conservation Bioacoustics, Cornell University, Ithaca, NY, USA

2. Freshwater and Oceanic Science Unit of Research, University of Liege, Liege, Belgium

3. Present address: Marine Mammal Research, Aarhus University, Roskilde, Denmark

4. Center for Avian Population Studies, Cornell Lab of Ornithology, Cornell University, Ithaca, NY, USA

Reproducible analytical workflows are critical for ensuring effective collaboration and efficient scientific workflows in the context of passive acoustic monitoring. Given that we deal with large amounts of data and extensive analytical pipelines, it is essential that the steps taken to process and work with passive acoustic data are reproducible and replicable. Open-source platforms that rely on version control (see below) provide a pathway for enabling scientists to share their workflows, documenting changes, and making it easier to collaborate. This guide outlines the best scientific practices for reproducibility.

### Why is reproducibility important and what are the steps involved?

Reproducibility is often considered the gold-standard for scientific rigor (Essawy et al. 2020) and there is often a misconception that one needs to worry about reproducibility only after the data collection is completed. A few key reasons why reproducibility is important include enhancing collaborative learning and research, meeting publication mandates, meeting funding agencies mandates, testing methods and analysis pipelines, increasing efficiency and transparency in research and minimizing data loss and reducing risk. There are three key steps involved in ensuring reproducibility of your PAM project: (*1*) data storage and organization; (*2*) best coding and file naming practices and (*3*) version control (Alston and Rick 2021).

#### 1. Data storage and organization

To ensure the data and its organization allow for transparent and reproducible analysis, we recommend that raw data and associated information be stored and documented in an efficient manner. A few key considerations here include: (*1*) Backing up raw data prior to analysis or any subsequent outputs in a regular manner. (*2*) Data must be organized in folders that are easy to read and are self-explanatory. (*3*) Using standardized variable names for folders and sub-folders and files. (*4*) Using appropriate data formats for the project in question and (*5*) Robustly documented metadata to explain the organization of data for you and others interested in working on the project.

An example of data organization include:

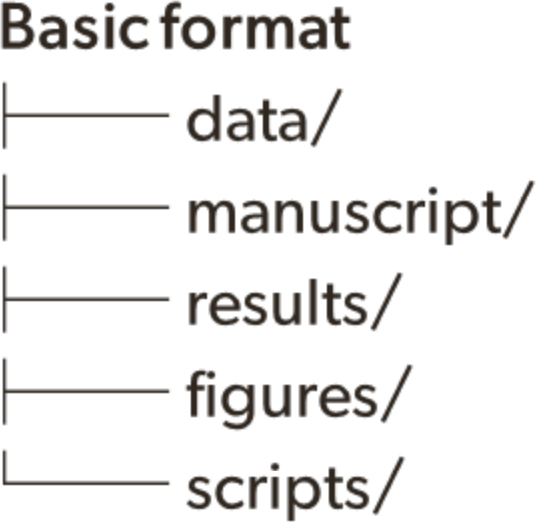

Here, all the raw audio data is stored in the data/ folder, any written reports go in the manuscript/ folder, while all subsequent analysis of audio data would be placed in the results/ folder. Further, figures and scripts would be organized in separate folders. Other examples of data storage and organization exist and for the sake of this guide, we provide a basic example above (Cooper & Hsing 2017).

#### 2. Best coding and file naming practices

For effective analysis of acoustic data, practicing a few important steps can help ensure your analysis is reproducible. These include reducing or removing spaces between names (e.g. spatial data vs. spatial-data; the use of hyphenation helps remove space here), providing content appropriate names for readability, use of lowercase letters and using numbers to illustrate chronological sequence of your scripts or files (e.g. 01_exploratory-analysis.Rmd; 02_statistical-models.Rmd). In a few passive acoustic projects, it is common for the sampling rate, as well as the date and time, to be included in folder or file names (e.g., ProjetGalapagos_site02_20240205_ 130000.wav for a file recorded at 1 PM on February 5). It is important to use a clear date format (in the example above, YYYYMMDD), as not all countries follow the same convention, and this should be explicitly stated. In addition, the use of local time or UTC should be clearly identified and is sometimes indicated by adding “_z” to the file name. In all cases, these conventions should be clearly explained in the accompanying documentation.

Besides naming your files and scripts and following appropriate naming conventions, it is also important to follow effective naming conventions for names of variables within a given file. For example, within a particular .csv, it is recommended that data is stored in a tidy format (variables in columns and observations in rows). Removing spaces in variable names also reduces any errors during processing and makes it easier to script the same. A practice we recommend is either using underscores, camelCase or snake-case to name your variables and following the same practice across all the files. For example, aru_sampling_time here refers to amount of time sampled while the passive acoustic monitor was recording. The same variable can be named as aruSamplingTime (referred to as using camelCase) or as aru-sampling-time (snake-case). Please note that using underscores is a version of snake-case. Please also ensure that within any given column in a file, the data structure is consistent. For instance, if the data structure consists of numbers or text or is alphanumeric in nature, please ensure that it remains consistent within the file and across files.

#### 3. Version control

Version control/version control systems (Blischak et al. 2016) allow you to essentially track and document the entire history of changes within a file, a script, or a project. One can easily tag a particular version and revisit their changes if needed. The introduction of version control has helped facilitate collaborations by making contributions transparent (Bryan, 2018).

Git is a version control software that allows you to track changes to your files over time. One can work with this software and interface with GitHub, a website for code and data (note: GitHub is not meant to store large amounts of data). GitHub is not only an archive but also a location where you can access files and data shared by collaborators. To enhance reproducible scientific practices, the use of version control via GitHub allows us to streamline our analysis and reduce time spent sifting through past work. In addition, errors that creep into the workflow and pipelines can be easily navigated and addressed due to version control.

##### Open-access materials

https://github.com/vjjan91/reproducibility-in-science (Here, we provide open-access materials to learn the basics of reproducible scientific practices and using GitHub).

### Conclusion

With the advent of larger passive acoustic datasets and rigorous deep learning pipelines being used to analyze data, following reproducible scientific practices allows scientists and practitioners to accelerate science and support conservation practice.

## Notes

### Competing Interest Statement

The authors have declared no competing interest.

### Summary of Updates

There was a mistake in the order of the authors enter in bioRxiv (the PDF has the correct order). Specifically, Cristian Perez-Granados is the last author not the second author. The order of authors is the only change in this revision.

